# mChIP-seq for Multiplex and Multifactorial Epigenomic Profiling Uncovers Cancer-specific Histone Features in Cellular and Circulating Nucleosomes

**DOI:** 10.64898/2026.04.27.721226

**Authors:** Changbin Sun, Qinkai Zhang, Jianli Yan, Xilin Wang, Chunxiao Zhang, Yujie Li, Jie Li, Wei Xu

## Abstract

Epigenomic profiling facilitates access to investigate regulatory roles of histone marks in a type-specific cell, and serves as a critical path for discovering noninvasive epigenetic models in cell-free nucleosomes. Here, we present mChIP-seq, an epigenomic profiling technology that is compatible with both cell and cell-free samples for synchronously profiling multifactorial epigenetic landscapes on multiple samples. Combining sample indexing in a single reaction with a pool-and-split strategy for immunoprecipitation, mChIP-seq enhances efficiency and reduces cost. Using mChIP-seq, we profiled H2A.Z and 10 histone modifications in cell lines representing 9 cancer types. Integrative analyses further revealed an atypical association of H2A.Z and H3K4me3 at promoter regions in cancer. Based on mChIP-seq, we developed cf-mChIP-seq for circulating nucleosomes, which requires as little as 25 μl of plasma per profile. Profiling 38 plasma samples for H2A.Z, H3K4me3, H3K27ac, and H3K9me3 with cf-mChIP-seq revealed distinct histone mark-associated cfDNA fragment patterns in breast cancer versus healthy control, highlighting the potential of cf-mChIP-seq to expand liquid biopsy methodologies. These results demonstrate that mChIP-seq is a widely applicable technology for large-scale epigenomic profiling of nucleosomes in cellular or cell-free forms.

## Introduction

The nucleosome, the basic repeating unit of chromatin in eukaryotes, is a DNA-histone complex consisting of approximately 145–147 base pairs of DNA wrapped around a histone octamer containing two copies of each of the four core histone proteins (H2A, H2B, H3, and H4) (1). Histone marks in nucleosomes, including histone post-translational modifications (PTMs) catalyzed by specific enzymes and histone variants, are dynamically distributed across the genome. These epigenetic signatures orchestrate chromatin state patterns to regulate spatiotemporal gene expression, determining cell fates (1–4). During cell death, fragmented chromatin is released into extracellular fluids (e.g., blood or urine), predominantly as cell-free nucleosomes (cf-nucleosomes) that preserve their original histone modification landscapes (5). Emerging evidence from epigenomic profiling studies in blood reveals that histone marks within cf-nucleosomes encode cell-type-specific gene expression signatures and fragmentomic patterns, offering opportunities to interrogate disease pathogenesis non-invasively (5–8). Therefore, comprehensive profiling of histone marks not only advances our understanding of their roles in cellular regulation but also serves as a critical path for discovering noninvasive epigenetic biomarkers or models in cf-nucleosomes with potential applications through liquid biopsy in multiple clinical settings.

ChIP-seq (chromatin immunoprecipitation followed by sequencing), first established in 2007, remains as the gold standard for epigenomic profiling of histone marks (4, 9–11). However, standard ChIP-seq necessitates substantial starting material, rendering it impractical for analyzing rare cell populations or low-abundance samples (10, 12). To address this limitation, several ChIP-seq derivatives have been developed by optimizing immunoprecipitation and sequencing library preparation, such as FARP-ChIP-seq (13) by adding carriers and ChIPmentation (14) by simplifying steps for library preparation. While these advancements improve sensitivity for low-input samples, their reliance on individual immunoprecipitation reaction inherently restricts processing throughput. To enable high-throughput epigenomic profiling, the multiplexing of barcoded samples in the same ChIP, which can be used as a strategy to reduce sample input as well, has been implemented in methods like iChIP (15), Bar-ChIP (16), Mint-ChIP (17), RELACS (18). and MINUTE-ChIP (19). Although these methods enable pooling multiple barcoded samples for joint processing to minimize technical variation while allowing to profile multiple epitopes from the same pool, current cell-free ChIP-seq (cfChIP-seq) for epigenomic profiling of cf-nucleosomes in liquid biopsy samples still face the two limitations: substantial starting materials (1ml or more of plasma per profile) and individual immunoprecipitation with low throughput (5, 7), underscoring the demand for technologies to reduce input requirements and improve processing efficiency.

Histone H2A.Z is a highly conserved H2A variant associated with many biological processes, such as transcriptional control, DNA repair and replication, and regulation of DNA accessibility and centromeric heterochromatin (20–23). Increased expression of H2A.Z has been observed in cancers including melanoma (24), breast (25), prostate(26), liver (27) cancers, etc., and is associated with poor prognosis (28, 29). However, there is still a lack of data to comprehensively characterize the epigenetic landscapes of H2A.Z to facilitate our understanding on the regulatory role of H2A.Z in cancer and investigate its features in plasma from patients with cancer.

Here we established an easy-to-use method for high-throughput and cost-effective epigenomic profiling, named mChIP-seq (**<u>m</u>**ultiplexed **<u>Ch</u>**romatin **I**mmuno**<u>P</u>**recipitation followed by **<u>seq</u>**uencing), based on the tailing and single-strand ligation method we developed (30) to label samples in a single reaction. We validated that mChIP-seq can stably and reproducibly profile various chromatin proteins for multiple samples in parallel with the advantages of high throughput, low-input requirement, and low cost. By using mChIP-seq, we systematically investigated epigenomic characteristics and associations of H2A.Z with 10 key histone modifications (H3K4me1, H3K4me2, H3K4me3, H3K9me3, H3K18ac, H3K27ac, H3K27me3, H3K36me1, H3K36me2, and H3K36me3) for 24 cancer cell lines across 9 cancer types, and revealed an atypical association of H2A.Z and H3K4me3 at promoter regions in cancer cells. Finally, we demonstrated that cell-free mChIP-seq (cf-mChIP-seq) requires as little as 25 μl of plasma per profile, a nearly 40-fold reduction in starting material requirement compared with conventional cfChIP-seq (typically 1 000 μl of plasma per profile). Profiling 38 plasma samples with breast cancer and non-cancer controls for H2A.Z, H3K4me3, H3K27ac, and H3K9me3 with cf-mChIP-seq, we obtained 156 profiles, observed the atypical association of H2A.Z and H3K4me3 in plasma with breast cancer as well, and found the different cfDNA fragment patterns associated with different histone marks, highlighting the potential of cf-mChIP-seq for expanding the methodological repertoire of liquid biopsy.

## Results

### Design and validation of mChIP-seq

To establish an easy-to-use multiplexing method for epigenomic profiling, we designed mChIP-seq as shown in Figure 1a. In mChIP-seq, the tailing and ssDNA (single-stranded DNA) ligation method, which we developed for high-throughput R-loop profiling (30), was used to label the 3’ end of DNA in fragmented chromatin with sample-specific barcodes. Then, the labeled samples were pooled and split for immunoprecipitation with relevant antibodies. After immunoprecipitation and ChIPed DNA purification, steps of extension, second adapter ligation, and indexing PCR were performed to prepare the sequencing library (30). Compared with other reported multiplexing methods using dsDNA (double-stranded DNA) adapter for sample indexing, the tailing and ssDNA ligation method in mChIP-seq substantially reduces the number of steps required for sample barcoding from two or three to one with high ligation efficiency (30) (Table S1, Supporting Information). Furthermore, mChIP-seq is designed with full consideration for the habits of users who are familiar to standard ChIP-seq by retaining the main operational steps in ChIP-seq protocol, which endows it with high adaptability and ease of use (Figure 1a). Multiplexed sample processing with multi-target immunoprecipitation enables high-throughput histone modification profiling of mChIP-seq, integrating advantages of low-input requirement (∼3,000 cells per target) and substantial cost reduction, such as 3-16 times cheaper in cost per library than typical ChIP-seq when 24 samples were pooled for targets from 1 to 12 (Figure S1a, Supporting Information).

**Figure 1.**
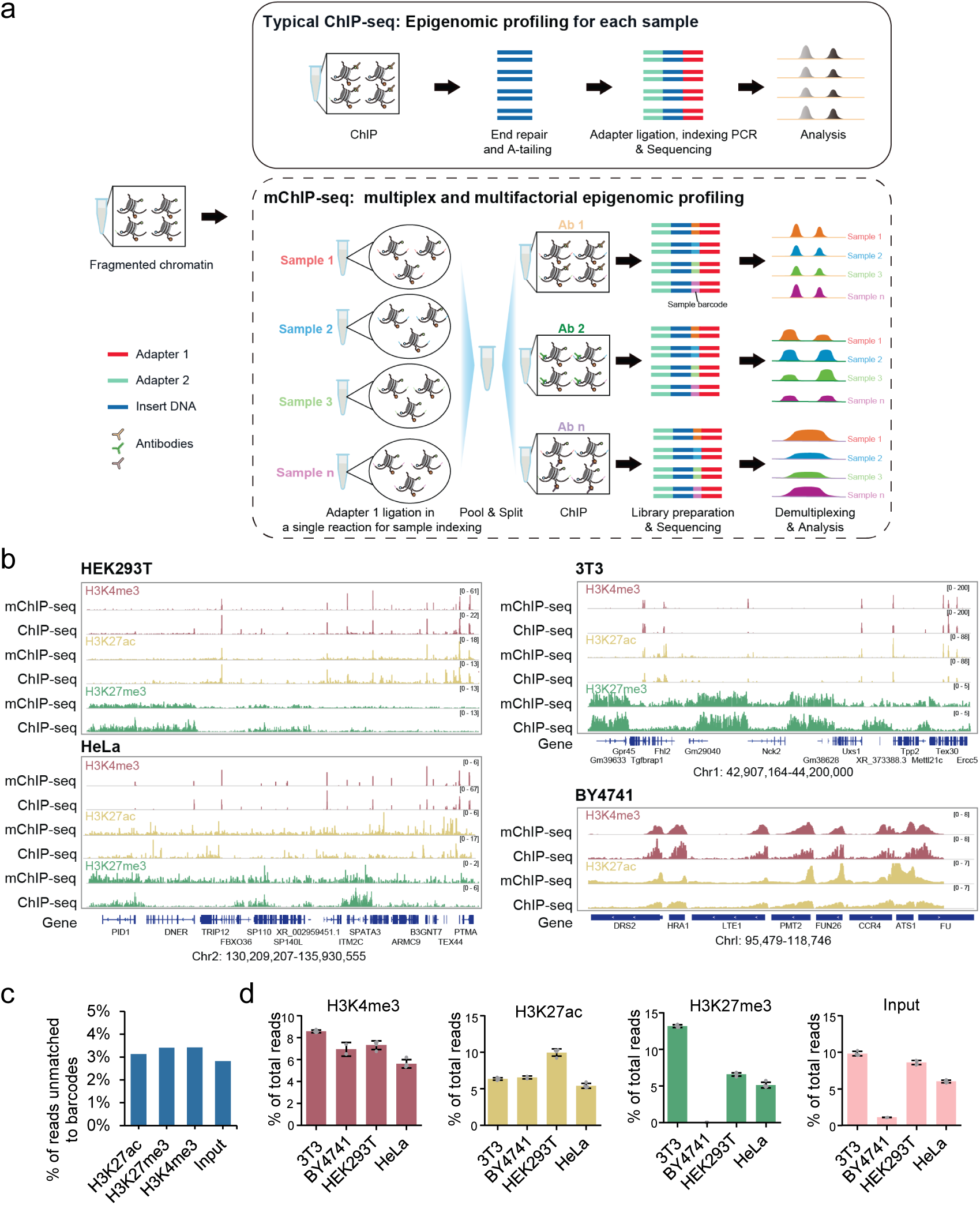
Design and validation of mChIP-seq for epigenomic profiling. a) Schematic diagram of the mChIP-seq and typical ChIP workflows. b) Genome browser tracks showing H3K4me3, H3K27ac, and H3K27me3 profiles derived by mChIP-seq for human cell lines HEK293T and HeLa, mouse cell line 3T3, and budding yeast strain BY4741, of which four replicates were indexed, pooled and split for the three modifications. Average signals of the replicates were represented. For comparison, ChIP-seq data from public database were shown. c) Bar plot showing percentage of reads unmatched to barcodes in the mChIP-seq data of the three modifications and Input. d) Bar plots depicting percentage of reads assigned to samples in the mChIP-seq data (n = 4).

We initially optimized the tailing and ssDNA ligation reaction for sample indexing, which is the key step in mChIP-seq. Given that detergents such as SDS and/or Triton X-100, which are typically present in sonication buffer to facilitate cell lysis for chromatin fragmentation, probably influence the ligation efficiency, we tested ligation reactions in buffers with different concentrations of SDS and Triton X-100. Our experiments demonstrated that when the SDS concentration was reduced to 0.05% in the presence of 1% Triton X-100, the inhibitory effect of SDS on ligation was remarkably mitigated (Figure S1b, Supporting Information). Under this optimized ligation condition, we assessed that about 18% of DNA 3’ ends in fragmented chromatin were barcoded without obvious GC bias (Figure S1c-e), and successfully profiled the histone modification H3K4me1 in budding yeast by ligating the first adapter to chromatin before immunoprecipitation (Figure S1f, Supporting Information). The data showed that the enriched signals were highly consistent between mChIP-seq and typical ChIP-seq (Figure S1g, Supporting Information).

Next, we validated mChIP-seq by pooling human cell lines HEK293T and HeLa, mouse cell line 3T3, and budding yeast strain BY4741 for histone modifications H3K27ac, H3K27me3, and H3K4me3 (Figure 1b). Demultiplexing results showed that more than 95% of the reads were assigned to one of the samples (Figure 1c). The overall alignment rate of each sample was high, averaging over 95%, while less than 0.5% of reads align specifically cross-species (Figure S2a, Supporting Information), indicating that the stop reaction effectively inhibited potential cross-contamination during the processes for sample mixing and immunoprecipitation. Additionally, only a quite small proportion (∼0.04%) of reads in the H3K27me3 data were assigned to the yeast strain BY4741 (Figure 1d). Budding yeast lacks the H3K27me3 modification in the genome (31), demonstrating the high specificity of mChIP-seq. Compared to standard ChIP-seq, the enriched signals identified by mChIP-seq were well-correlated with those of ChIP-seq (Figure S2b-e, Supporting Information).

To further evaluate the reproducibility and specificity of mChIP-seq for tissues, we profiled H3K4me3 and H3K27ac in liver and pancreas tissues from two pig breeds, each sample indexed with three barcodes (Figure S3, Supporting Information). At the genome-wide, correlation analysis demonstrated high consistency of signals among replicates (Figure S4, Supporting Information). To demonstrate the strengths of mChIP-seq on the comparative analysis of histone modification levels across different biological conditions, we generated H3K27ac ChIP-seq data using the same materials. Between the two methods, about 80% of peaks overlap, similar to the percentage of peaks overlapping among replicates of each method (Figure S5a, Supporting Information). The correlation analysis demonstrated high consistency in signal patterns between mChIP-seq and ChIP-seq profiles (r range from 0.97 to 0.99) (Figure S5b, Supporting Information). Differential analysis showed that around 90% of peaks were commonly assigned as differential enrichment sites (DESs, log_2_(fold change) ≥ 1 or ≤ -1 and FDR ≤ 0.001), including DESs associated with well-known liver-specific genes such as *SPERPINC1* and *SPP2* and pancreas-specific gene such as *PNLIP* and *PNLIPRP1* (Figure S5c-f, Supporting Information). In addition, peaks assigned as DESs only in one method are those with relatively low enrichment scores (Figure S5g, Supporting Information). We also successfully profiled RNA polymerase II (Pol II) in these tissues with relatively higher materials (See Experimental Section), demonstrating mChIP-seq is compatible with DNA binding protein profiling (Figure S6, Supporting Information). Taken together, our results demonstrated that mChIP-seq can be used to multiplex chromatin landscape profiling with similar performance as traditional ChIP-seq for quantitative comparison.

### Profiling of pan-cancer histone mark atlas with mChIP-seq

The histone variant H2A.Z has been demonstrated to play an important role in tumor growth and chemoresistance, but its mechanisms in transcriptional regulation remain largely elusive (20, 28). We applied mChIP-seq to profile H2A.Z and 10 well-studied modifications of H3 histone (H3K4me1, H3K4me2, H3K4me3, H3K9me3, H3K18ac, H3K27ac, H3K27me3, H3K36me1, H3K36me2, and H3K36me3) in 24 cancer cell lines representing a diverse array of tissue types, including blood, breast, colon, central nervous system, kidney, lung, ovary, prostate, and skin (32) (Figure 2 a,b). These 24 cancer cell lines were chosen from the NCI-60 panel, in which the K562 cell line, an Encyclopedia of DNA Elements (ENCODE) project tier 1 cell line and one of the most commonly used human myelogenous leukemia cell was included to monitor the data quality (32, 33). We totally generated 576 libraries from 22 chromatin immunoprecipitation assays for these marks and Input without immunoprecipitation (Table S2). On average, approximately 99% of the reads were successfully assigned to one of the barcodes (Figure S7a, Supporting Information), and about 86% of the mapped reads were uniquely aligned to the genome (Figure S7b, Supporting Information). Consistent with previous reports (2, 3, 34), the signals of H2A.Z, H3K4 methylations (H3K4me1, H3K4me2, and H3K4me3), H3K27ac, and H3K18ac, which are known to be associated with promoters and/or enhancers, exhibited a bimodal distribution surrounding the transcription start sites (Figure 2c,d). The signals of H3K36me3, which are typically associated with transcribed regions, showed elevated enrichment in the gene body (Figure 2c,d). In contrast, the signals of H3K36me1, H3K36me2, H3K27me3, and H3K9me3, marks related to repressed states and/or repetitive states (35), were relatively diminished in the gene body (Figure 2c,d).

**Figure 2.**
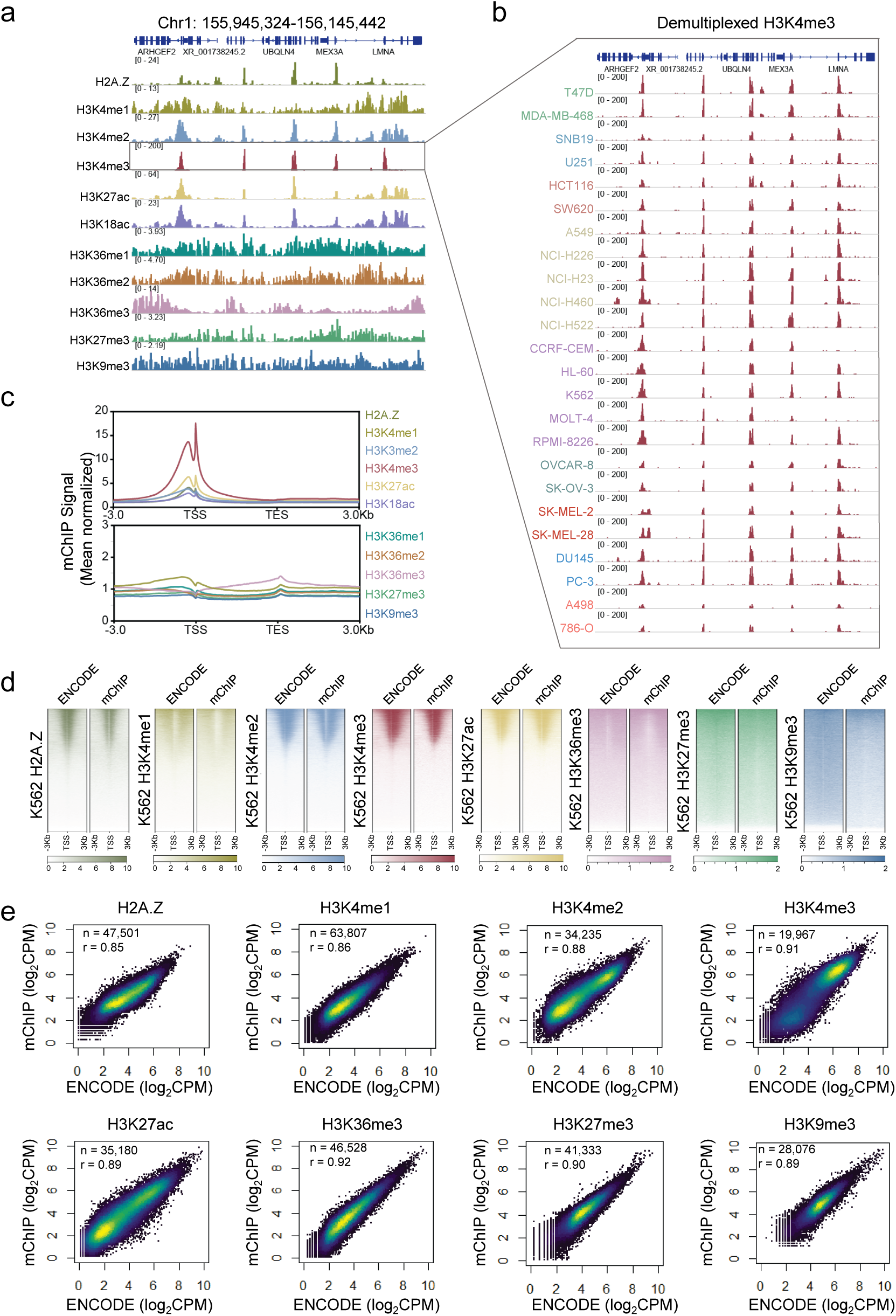
Pan-cancer multifactorial histone mark atlas profiling with mChIP-seq. a) Genome browser tracks showing profiles of histone variant H2A.Z and 10 histone modifications derived by mChIP-seq for 24 cancer cells across 9 cancer types without de-multiplexing. 50 million raw reads were sampled and used for each histone mark. b) Representative genome browser tracks of H3K4me3 after de-multiplexing. c) Plots showing normalized signal intensities over transcripts. Flanking regions are 3 kb upstream of TSSs and 3 kb downstream of the TESs. d) Heatmap showing signal intensities around TSSs for K562 histone marks derived by mChIP-seq and ChIP-seq from ENCODE. Signal intensities were sorted by ENCODE data. e) Heatscatter plots showing the correlations in peaks between mChIP-seq and ENCODE. n: number of peaks; r: Pearson correlation coefficient.

To evaluate the data quality of mChIP-seq upon expanding the size of samples to 24 and histone marks to 11, we compared K562 mChIP-seq data sets, including H2A.Z, H3K4me1, H3K4me2, H3K4me3, H3K9me3, H3K27ac, H3K27me3, and H3K36me3, to available analogous ENCODE data generated by standard ChIP-seq (33). The genomic tracks of these histone marks and signals surrounding the TSSs showed similar distributions of enrichment in profiles with mChIP-seq relative to the ENCODE datasets (Figure S8a, Supporting Information). In addition, the identified enrichment sites demonstrated substantial overlap (Figure S8b, Supporting Information) and exhibited robust correlations between the respective datasets, with Spearman correlation coefficients of 0.85, 0.86, 0.88, 0.91, 0.89, 0.92, 0.90, and 0.89 for H2A.Z, H3K4me1, H3K4me2, H3K4me3, H3K9me3, H3K27ac, H3K27me3, and H3K36me3, respectively (Figure 2e). Furthermore, similar signal-to-noise ratios as demonstrated by FRiP (fraction of reads in peaks) were observed between mChIP-seq and ENCODE data sets (Figure S8c). These results provided compelling evidence of the high reproducibility and reliability of mChIP-seq for the parallel profiling of up to 24 samples and 11 histone marks.

### Epigenomic characterization of H2A.Z in cancer cells

The data sets generated with mChIP-seq enabled comprehensive epigenomic characterization of H2A.Z in these cancer cells. Three replicates were performed for H2A.Z using antibodies from two manufacturers (Table S2). Enriched regions identified in all replicates of each cell line were assigned as reproducible enrichment sites for the following analysis. Based on these data sets, we totally identified 188,464 enrichment sites of H2A.Z, with an average of 19,655 H2A.Z enrichment sites per sample (Figure 3a; FigureS9, Supporting Information). Annotation analysis demonstrated that the majority of H2A.Z enrichment sites were mapped to genomic loci associated with promoter, intron, and distal intergenic region. On average, each cell line exhibited 51.73% enrichment sits at promoter, 19.72% at intron, and 24.31% at distal intergenic region. H2A.Z levels exhibited a relatively higher correlation with those of promoter marks (H3K4me3 and H3K4me2) and active marks (H3K27ac and H3K18ac) at promoter sites, as well as a higher correlation with levels of enhancer associated mark H3K4me1 and active marks (H3K27ac and H3K18ac) at non-promoter sites (Figure 3b,c; Figure S9c, Supporting Information). In addition, H2A.Z signals were positively correlated with chromatin accessibility assessed by ATAC-seq (Figure S9d, Supporting Information), which is in line with previous finding that H2A.Z deposition may enhance nucleosome mobility and DNA accessibility (23).

**Figure 3.**
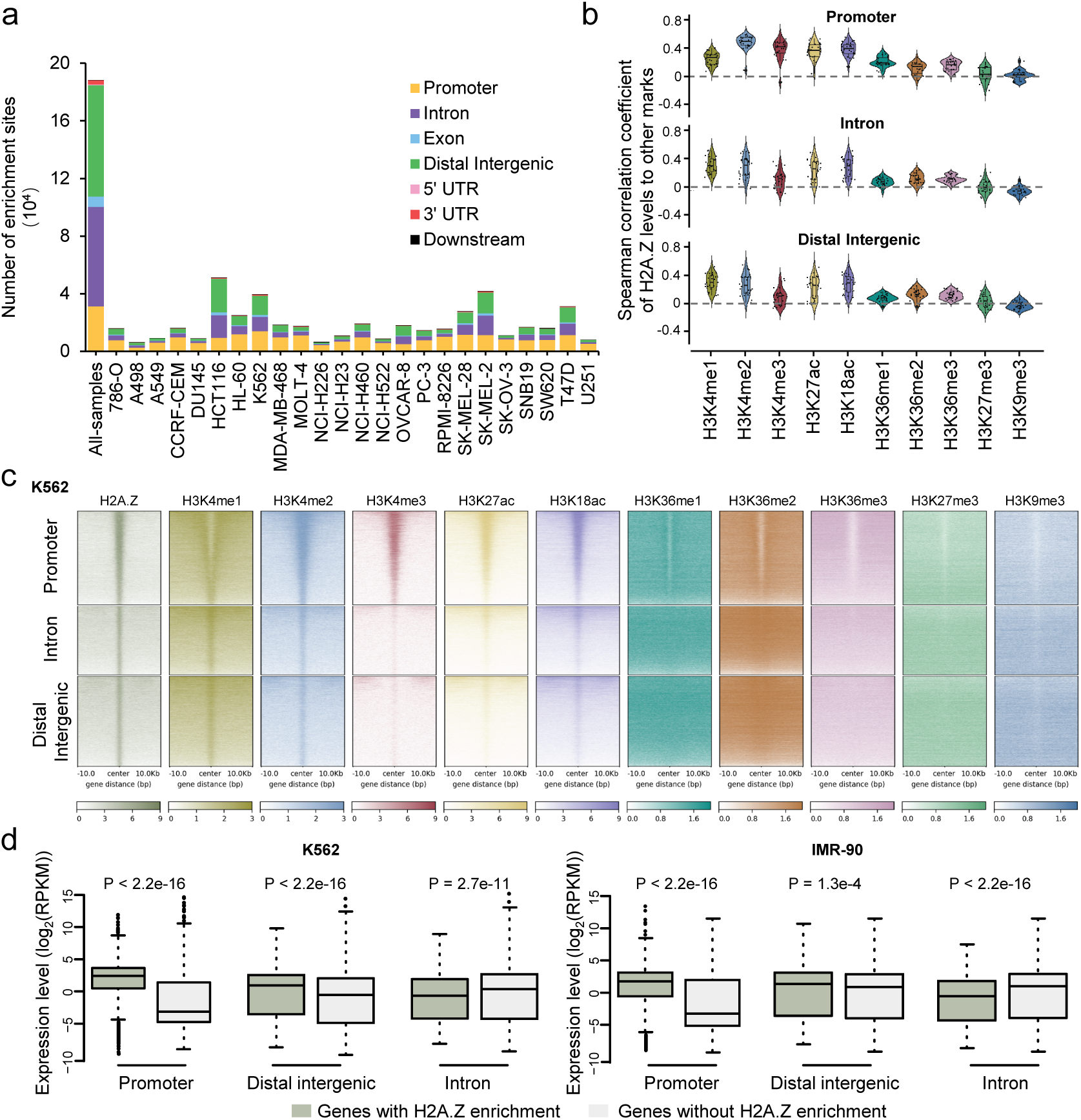
Epigenome characterization of H2A.Z in cancer cells. a) Number of H2A.Z enrichment sites obtained from mChIP-seq data. All-samples represents the merged peak set from all samples. Color indicates the type of the genomic region overlapped by the enrichment sites. UTR: untranslated region. b) Violin plots showing correlations of H2A.Z to other histone marks for each cell line in the H2A.Z enrichment sites localized in promoter, intron, and distal intergenic regions. c) Representative Heatmap showing signal intensities of 10 histone modifications around H2A.Z enrichment sites localized in promoter, intron, and distal intergenic regions identified in K562. Signal intensities were sorted by H2A.Z data. d) Boxplots comparing expression level of genes with enrichment H2A.Z with genes without H2A.Z enrichment in cell lines K562 (left) and IMR-90 (right). All P values were determined by Wilcoxon rank-sum tests.

To elucidate the influence of H2A.Z deposition on gene expression, RNA-seq data of the corresponding cancer cell lines (Table S3, Supporting Information) were integrated for the correlation analysis. The results showed strong positive correlation between H2A.Z and gene expression levels at promoter sites (Figure 3d; Figure S10a, Supporting Information). Additionally, genes with H2A.Z enrichment in distal intergenic regions exhibited higher expression levels compared with those without such enrichment, suggesting that H2A.Z plays a transcriptional activation role at these regulatory elements (Figure 3d; Figure S10b, Supporting Information). In contrast, the role of H2A.Z in transcriptional repression were also observed for genes with H2A.Z enrichment at intron sites (Figure 3d; Figure S10c, Supporting Information), though levels of H2A.Z positively correlated with levels of enhancer associated mark H3K4me1 and active marks (H3K27ac and H3K18ac) at these H2A.Z intron sites (Figure 3b). These results indicated that the functional impact of H2A.Z on gene expression regulation is dependent on its genomic location. Notably, similar transcriptional regulatory characteristics of H2A.Z were also found in non-cancerous cell lines based on public datasets (Figure 3d; Figure S11, Supporting Information). Collectively, these results demonstrated that H2A.Z was predominantly deposited at promoters and putative enhancer regions, supporting its crucial roles in transcriptional regulation by modulating promoter and enhancer activity (36, 37).

### Epigenetic dysregulation of H2A.Z in cancer cells

Overexpression of H2A.Z has been observed in various cancer types, yet its influence on gene expression remains unclear (20). To investigate the impact of H2A.Z overexpression on its genomic localization, we compared H2A.Z expression levels (*H2AZ1* and *H2AZ2*) of cancer cell lines with non-cancerous cell lines (Table S3). In general, cancer cell lines exhibited higher expression levels of *H2AZ1* and *H2AZ2* than non-cancerous cell lines (Figure 4a; Figure S12a, Supporting Information). For instance, HCT116 is one of the cell lines with the highest *H2AZ1* expression (Figure 4a). Notably, a slightly negative correlation of signals between H2A.Z and H3K4me3 at promoter sites in HCT116 according to the correlation analysis was shown in Figure 3b, corroborated by independent public data (Figure S12b, Supporting Information). Considering the strong correlation between H2A.Z levels at promoters and gene expression (Figure S10 and S11, Supporting Information), we focused on the effect of H2A.Z overexpression on its deposition at promoter regions in cancer cells by analyzing its associations with the promoter mark H3K4me3. To determine whether non-cancerous cells exhibit distinct distributions of H2A.Z, we therefore performed correlation analyses between H3K4me3 and H2A.Z signals at promoter H2A.Z enrichment sites across downloaded data sets for comparison (Table S3, Supporting Information). We also calculated the correlations of H3K4me3 to H3K27ac and H3K27me3, two main marks associated with active and repressed genes, respectively. Our results demonstrated that most cancer cell lines showed positive correlations between levels of H2A.Z and H3K4me3 at promoter H2A.Z enrichment sites. However, the correlation coefficients in cancer cell lines were more variable and significantly lower than those in non-cancerous cell lines (Figure 4b). In contrast, the associations between H3K4me3 and H3K27ac and H3K4me3 and H3K27me3 were nearly equivalent between the two cell types, suggesting that H2A.Z accumulation might be dysregulated at promoter sites in some cancer cell lines.

**Figure 4.**
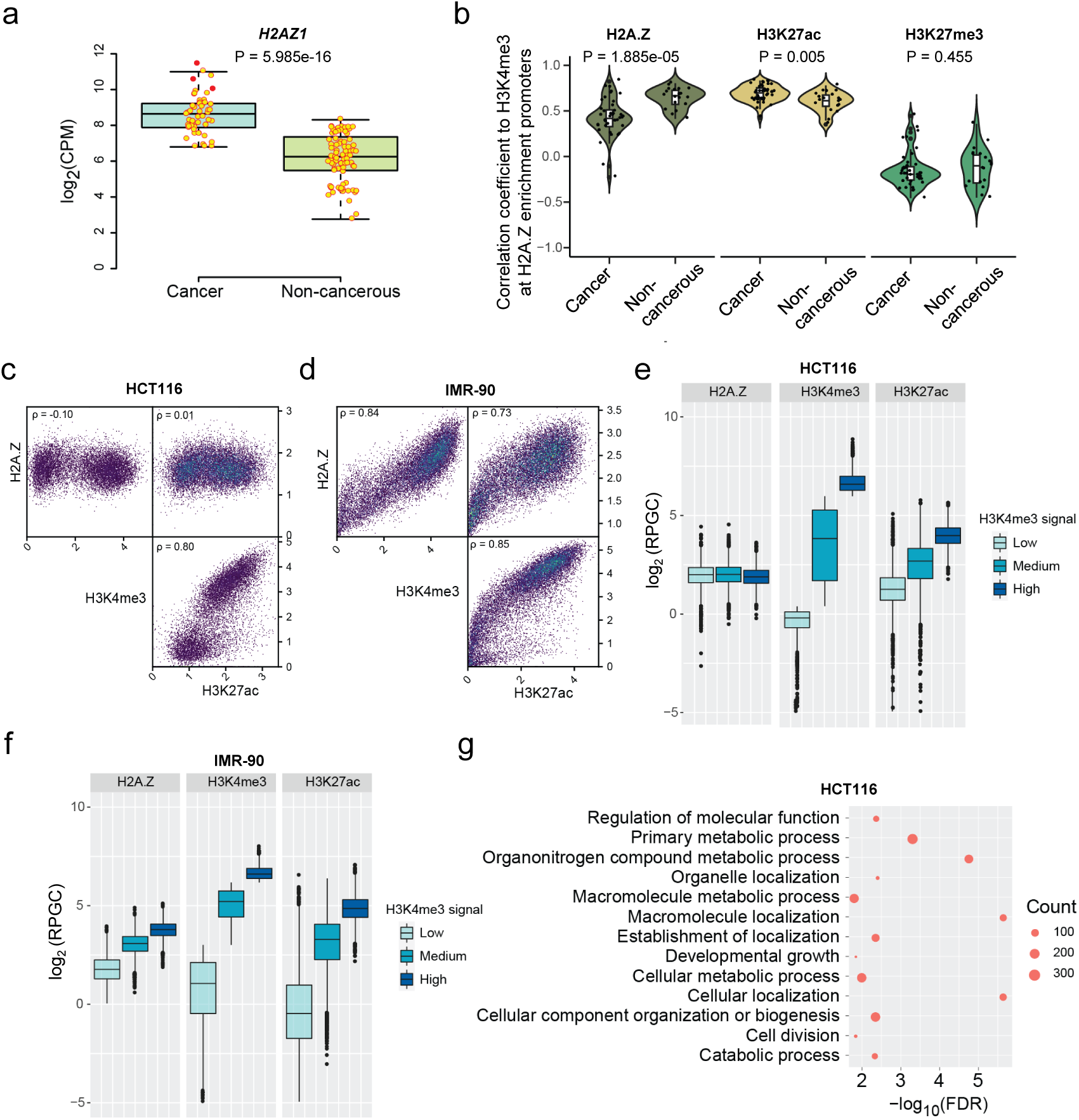
Epigenetic dysregulation of H2A.Z in cancer cells. a) Boxplot comparing expression of *H2AZ1* in cancer cell samples (n = 56) with non-cancerous cell samples (n = 65). P values were calculated using Mann-Whitney U tests. b) Violin plots comparing correlations of H3K4me3 to H2A.Z and H3K27ac in cancer cell lines (n = 41) with non-cancerous cell lines (n = 19) at promoter H2A.Z enrichment sites. P values were calculated using Mann-Whitney U tests. a) and d) Pairwise heatscatter plots showing the correlations of signal among H2A.Z, H3K4me3, and H3K27ac at promoter H2A.Z enrichment sites in HCT116 (**c**) and IMR-90 (**d**). ρ: Spearman correlation coefficient. e) and f) Boxplots showing signal distribution of H2A.Z at promoter H2A.Z enrichment sites with different H3K4me3 intensities in HCT116 (e) and IMR-90 (f). g) Results of GO biological process enrichment analysis for expressed genes (RPKM > 1) with H2A.Z (RPGC > 1) and without H3K4me3 (RPGC < 1) signals at promoter in HCT116. Fifteen terms with the lowest FDR were presented.

Interestingly, we observed comparable H2A.Z intensity across promoter H2A.Z enrichment sites with varying H3K4me3 intensities in HCT116 (Figure 4c), unlike the non-cancerous cell line IMR-90 (Figure 4d). By proportionally dividing the promoter H2A.Z enrichment sites into three groups based on the intensity of H3K4me3 (High, Medium, and Low), we found that H2A.Z showed comparable signal levels between the Medium and Low groups and even slightly lower in the High group in the cancer cell line HCT116 (Figure 4e). Differently, in the non-cancerous cell line IMR-90, the signal intensity of H2A.Z generally corresponded to that of H3K4me3 (Figure 4f). Similar patterns of H2A.Z dysregulation were observed in some other cancer cell lines, such as K562 and HepG2, unlike their non-cancerous counterparts, B cells and hepatocyte (Figure S12c-f). Together, these results suggest that H2A.Z overexpression in cancer cells may lead to an aberrant association of H2A.Z and H3K4me3 at promoters, potentially contributing to further epigenetic dysregulation of gene expression.

### The impact of abnormal H2A.Z accumulation on gene expression at promoter regions in cancer cells

To investigate the effects of dysregulated H2A.Z accumulation on gene expression at promoter regions in cancer cells, we performed a comparative expression analysis of genes with promoter H2A.Z enrichment, stratified into three groups based on the H3K4me3 signal intensity, between the cancer cell line HCT116 and the non-cancerous cell line IMR-90 (Figure 4 E and F). We observed that the highest signals of H3K4me3 coincided with the highest levels of gene expression (Figure S13a,b, Supporting Information), though the intensities of H2A.Z were nearly similar across promotor enrichment sites in HCT116 (Figure 4e). This finding suggested that the transcriptional activation function of H2A.Z at promoter sites might be largely dependent on its co-occurrence with H3K4me3 in cancer cells. We further compared the expression levels of genes (RPKM > 1) with H2A.Z (RPGC > 1) and without H3K4me3 (RPGC < 1) signals at the promoter. We observed that the expression levels of these genes were higher in cancer cells than non-cancerous cells (Figure S13c, Supporting Information), indicating the transcriptional activation function of H2A.Z might lead to dysregulated expression of genes with low H3K4me3 levels in cancer cells. Notably, Gene Ontology (GO) enrichment analysis demonstrated that these genes were enriched in metabolic process, biosynthetic process, and cell cycle regulation (Figure 4g; Figure S13 d-h, Supporting Information), which might partially explain the roles of H2A.Z overexpression in cancer cells (25, 28).

Unlike canonical histones H2A, the expression of the histone variant H2A.Z is independent of the replicative state, allowing its deposition throughout the cell cycle (20). SRCAP and EP400 are known chromatin remodelers in H2A.Z deposition, while ANP32E is an H2A.Z-specific chaperone capable of removing H2A.Z and promoting the establishment of H2A.Z-depleted chromatin loci (38–40). Dysregulation of these genes may also influence H2A.Z deposition in cancer cells. To investigate the relationship of these genes with H2A.Z, we compared their expression levels in cancer cells with non-cancerous cells. Generally, cancer cell lines exhibited higher expression levels of *SRCAP*, *EP400*, and *ANP32E* in comparison with non-cancerous cell lines (Figure S14a-c, Supporting Information). However, similar to non-cancerous cells, the expression levels of *SRCAP* and *EP400* negatively correlated with those of *H2AZ1* and *H2AZ2,* while the expression level of *ANP32E* positively correlated with those of *H2AZ1* and *H2AZ2* in cancer cells (Figure S14d,e). These results suggested a possible feedback regulatory mechanism that coordinate the expression of these genes to modulate H2A.Z deposition in the genome. Increasing evidence suggests that the dysregulation of *SRCAP*, *EP400*, and *ANP32E* expression is strongly associated with carcinogenesis (41, 42). Therefore, whether the abnormal deposition of H2A.Z in cancer cells is a consequence or a contributing factor to its overexpression requires further investigation. Together, these results demonstrated an abnormal accumulation of H2A.Z at promoter sites in cancer cells, which we speculate may contribute to the dysregulation of gene expression involved in critical biological processes such as cancer cell proliferation and viability.

### High-throughput and ultralow-input liquid biopsy epigenomic profiling with cf-mChIP-seq

Development of methodologies that are easier to implement on a large scale represents a crucial research direction in liquid biopsy (43). To address this need, we further improved mChIP-seq to enable the high-throughput and ultralow-input epigenomic profiling of cf-nucleosomes in plasma (Figure 5a). We adapted the pipeline of mChIP-seq for use with plasma samples and developed cf-mChIP-seq that allowed simultaneously profiling of multiple histone marks with starting materials as low as 25 μl of plasma per profile. Briefly, isolated plasma samples were directly indexed without DNA fragmentation. Indexed samples were subsequently mixed and pre-cleared with Protein A/G magnetic beads to remove potential native antibodies in plasma. Then, the pre-cleared mixture was split for ChIP with relevant antibodies followed by the rest library preparation steps as mChIP-seq protocol (Figure 5a, see Experimental Section for detail). As widespread DNA damage indicated in cfDNA (e.g., single-strand nicks and DNA overhangs) (44), cf-mChIP-seq that employs single-stranded adapter ligation without any DNA digestion or insertion are more faithfully preserved intactness of cfDNA fragments in the final libraries than conventional double-strand based libraries, enhancing the accuracy of cfDNA fragment size analysis.

**Figure 5.**
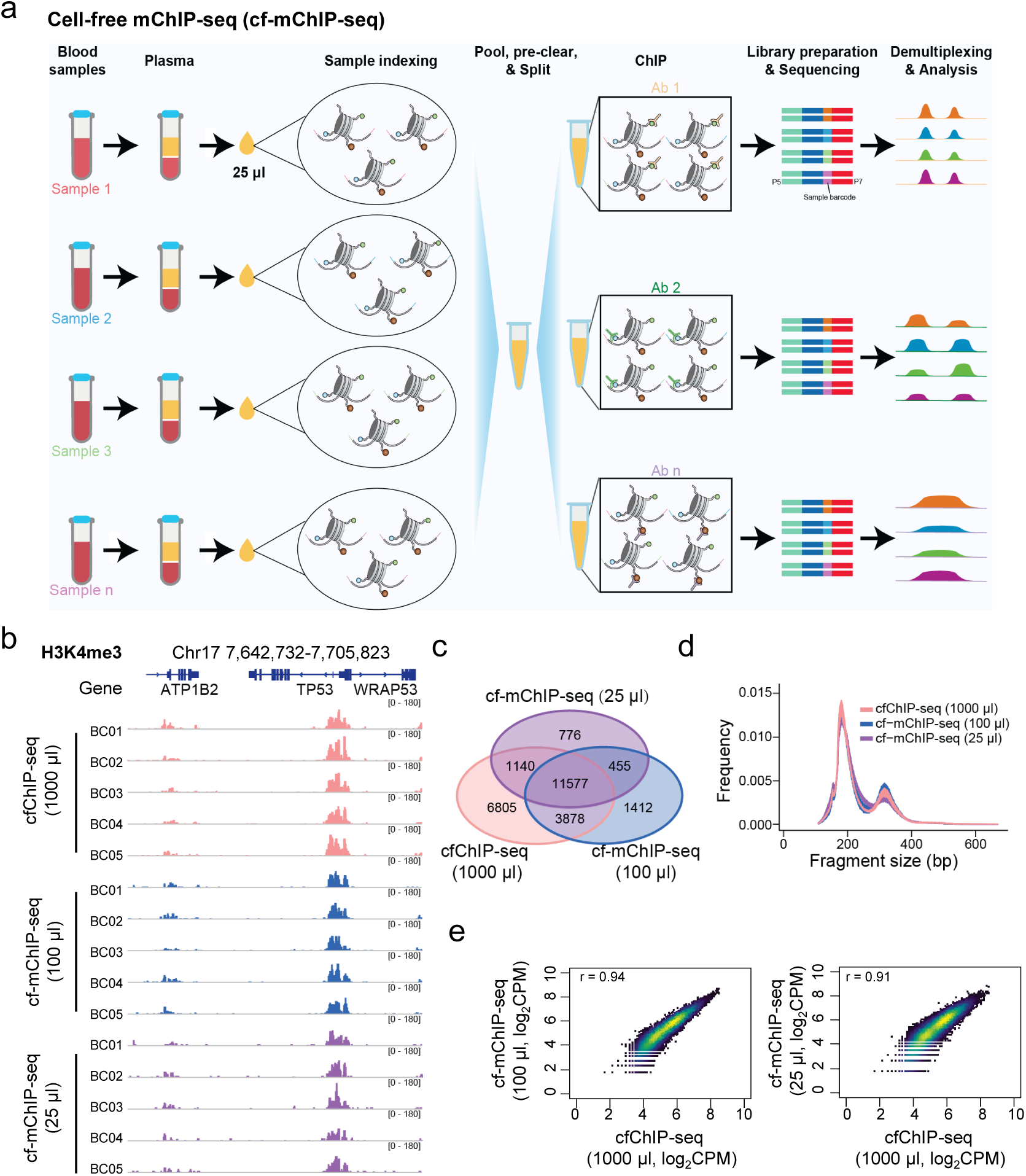
High-throughput and ultralow-input liquid biopsy epigenomic profiling with cf-mChIP-seq. a) Schematic overview of the cf-mChIP-seq. b) Genome browser tracks showing normalized signals of H3K4me3 mapped by cfChIP-seq (1000 μl), cf-mChIP-seq (100 μl), and cf-mChIP-seq (25 μl) for five plasma samples from patients with breast cancer. The number in the parentheses indicated the volumes of plasma used per profile. c) Venn diagram representing the number of H3K4me3 peaks shared among cfChIP-seq and cf-mChIP-seq with different input in BC05 sample. d) Size frequency distribution of sequenced fragments. n = 5 for each. e) Heatscatter plots showing the correlations in peaks between cf-mChIP-seq and cfChIP-seq data sets from BC05 sample. r: Pearson correlation coefficient. Peaks from cfChIP-seq were used for Pearson correlation analysis.

We benchmarked the performance of cf-mChIP-seq by using the antibody against H3K4me3 in plasma samples from five patients with breast cancer. Barcoded chromatin was prepared from two different volumes of input plasma (25 μl and 100 μl) as start materials for testing. Meanwhile, cfChIP-seq was simultaneously performed as described previously for comparison (7). As expected, strong H3K4me3 signals were observed around TSSs from both cfChIP-seq and cf-mChIP-seq profiles (Figure 5b). The genomic regions exhibiting significant signal enrichment in cf-mChIP-seq profiles demonstrated substantial overlap with corresponding profiles obtained through cfChIP-seq, even though dramatically reduced plasma may lose some weak signals (Figure 5c). Fragment size distributions of these data sets showed similar patterns, largely corresponding to DNA wrapped around mono- and di-nucleosomes as reported previously (5) (Figure 5d). Comparisons of signal intensities at enrichment sites confirmed high correlations between cf-mChIP-seq and cfChIP-seq profiles (Figure 5e). Thus, cf-mChIP-seq enables multiplexed epigenetic profiling of cf-nucleosomes from ultralow-input plasma sample.

### Epigenomic profiling of circulating nucleosomes in plasma from breast cancer patients using cf-mChIP-seq

We applied cf-mChIP-seq to profile H2A.Z, H3K4me3, H3K27ac, and H3K9me3 in 38 plasma samples (21 from breast cancer patients and 17 from healthy donors) and generated 156 cf-nucleosome epigenomic profiles (Figure S15a; Table S4, Supporting Information). Due to the lower input (25 μl per profile), cf-mChIP-seq yielded fewer usable reads per profile than typical cfChIP-seq (5, 7), with median values of 0.1 million in H2A.Z, 0.2 million in H3K4me3, 0.2 million in H3K27ac, and 2.9 million in H3K9me3 datasets, respectively (Table S4). However, the genomic distributions of cf-mChIP-seq signals around genes are consistent with those observed in cells (Figure S15b, Supporting Information). In total, 47,320, 58,614, 59,872, and 184,357 enrichment sites were identified for H2A.Z, H3K4me3, H3K27ac, and H3K9me3, respectively (Figure S15c, Supporting Information). Consistent with previous epigenetic study (5), dimensionality reduction by signal intensity in peaks demonstrated relatively lower variability of signals in peaks among healthy controls compared with cancer samples (Figure S15d, Supporting Information).

Based on our epigenomic profiling data sets in cells, breast cancer cell lines MDA-MB-468 and T47D showed lower correlations of H2A.Z and H3K4me3 at promoter sites than mammary epithelial cell (Figure 6a-c). As the circulating tumor DNA (ctDNA) refers to fragments of cfDNA that are released from tumor cells (45), we hypothesized that this aberrant feature of H2A.Z at promoter sites might occur in plasma from breast cancer patients. To evaluate this, we calculated correlation coefficients of H3K4me3 and H2A.Z and observed lower correlations in patients than those in healthy controls (p = 0.006, Wilcoxon rank-sum test; Figure 6d). Though the correlation coefficients of H3K4me3 and H3K27ac were more variable in patients than those of healthy controls, no significant difference between patients and healthy controls was observed (p = 0.531, Wilcoxon rank-sum test; Figure 6e). Notably, fragment size analysis revealed significantly shorter H2A.Z-associated cfDNA in cancer patients at enriched promoters (P < 0.0001, Wilcoxon rank-sum test), as previous reports that tumor-derived cfDNA are shorter than the non-tumor cfDNA (43) (Figure 6f). Further analysis demonstrated significant differences of median lengths among cfDNA fragments associated with different histone markers, though the fragment lengths are largely between 90bp and 450 bp, which corresponds to mono-(90-250 bp) and di-(251-450 bp) nucleosome-derived cfDNA fragments (Table S4). The median size of H3K4me3-associated cfDNA fragments were the longest, followed by H3K27ac, H2A.Z, and H3K9me3, and all histone mark-associated fragments were significantly longer than Input profiles (total cfDNA without immunoprecipitation) (P < 0.0001, paired t-test, Figure S16a, Supporting Information).

**Figure 6.**
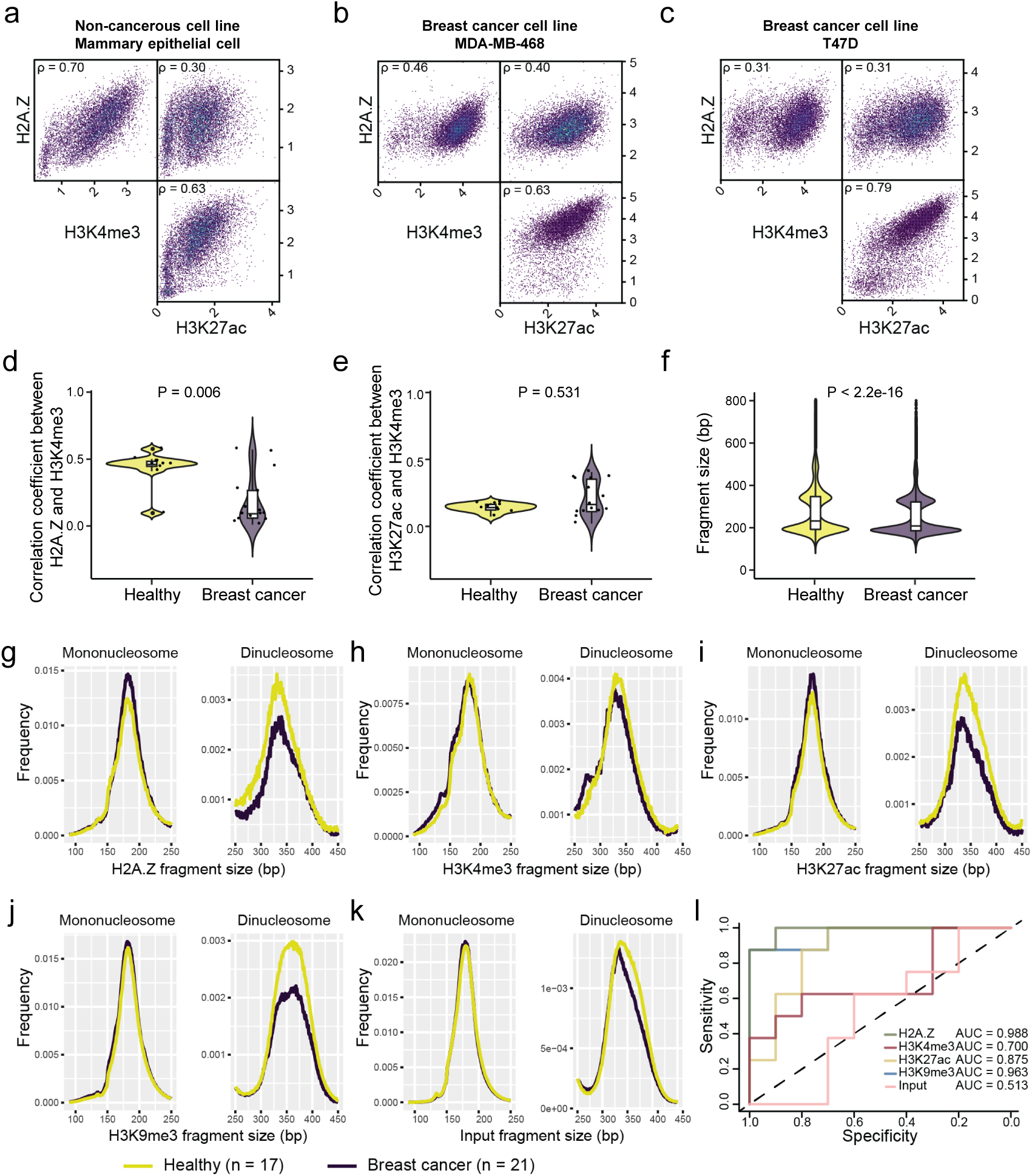
Epigenomic profiling of circulating nucleosomes in plasma from breast cancer patients using cf-mChIP-seq. a) to c) Pairwise heatscatter plots showing the correlations of signal among H2A.Z, H3K4me3, and H3K27ac at promoter H2A.Z enrichment sites in non-cancerous mammary epithelial cell (a), breast cancer cell lines MDA-MB-468 (b) and T47D (c), ρ: Spearman correlation coefficient. d) and e) Violin plot showing correlations of signals between H2A.Z and H3K4me3 (d) and between H2A.Z and H3K27ac (e) at promoter H2A.Z enrichment sites in plasma from healthy controls (n = 10) and breast cancer patients (n = 15). P value was determined by Wilcoxon rank-sum test. f) Violin plot comparing fragment size distribution of H2A.Z-associated cfDNA in H2A.Z enrichment promoters between no cancer and breast cancer plasma. P value was determined by Wilcoxon rank-sum test. g) to k) Line plots depicting the median fragment size frequency normalized by total fragment counts grouped by cancer status across the mono- and di-nucleosome fractions of cfDNA associated with H2A.Z (g), H3K4me3 (h), H3K27ac (i), H3K9me3 (j), and Input (k). l) ROC curves showing performances of classifiers using fragmentation patterns associated with different histone marks.

Next, we investigated the potential of cfDNA fragmentation patterns associated with histone marks to serve as cancer classifiers. The results showed that cfDNA fragments from breast cancer patients were shorter than those from healthy individuals (median sizes: cancer vs healthy—H2A.Z, 194.2 vs. 202.2 bp; H3K27ac, 200.3 vs. 228.0 bp; H3K4me3, 232.1 vs. 255.9 bp; H3K9me3, 189.0 vs. 197.4 bp; Input, 181.1 vs. 182.9 bp). Notably, significant differences (P < 0.001, Welch’s t-test) were observed for H2A.Z-, H3K27ac-, and H3K9me3-cfDNA, while H3K4me3-cfDNA showed weaker significance (P < 0.05), and Input displayed no significant difference (P > 0.05) (Figure S16b, Supporting Information). Focusing on fragments in the mono- and di-nucleosome fractions based on fragment size frequency and cumulative frequencies, distinct cfDNA size profiles were found between cancer and healthy samples. Specifically, cancer samples exhibited significantly reduced di-nucleosome fractions with H2A.Z, H3K27ac, and H3K9me3 (P < 0.001, Kolmogorov-Smirnov test) (Figure 6g-k; Figure S17a-e, Supporting Information). These differences were also obviously showed at the genome-wide level using the ratios of mono-nucleosome to di-nucleosome fragments in 5-megabase (Mb) windows (Figure S17f, Supporting Information). Using the genome-wide fragmentation patterns, we implemented a weighted elastic net machine learning model to create classifiers with these epigenetic profiles to test their performances for cancer detection. To this end, the same randomly selected half-samples were employed for training each histone mark and Input data using fivefold nested cross-validation and the left half-samples were used for validation. Results demonstrated high classification performances of fragmentation patterns associated with H2A.Z and H3K9me3, with an area under the curve (AUC) of 0.988 and 0.963, respectively (Figure 6l). Together, these results highlight proof of concept that cf-mChIP-seq can be implemented on a large scale with the potential for expanding the methodological repertoire of liquid biopsy.

## Discussion

In this study, we developed mChIP-seq, an easy-to-use method combining the pool-and-split strategy with sequencing to enable high-throughput and low-cost epigenomic profiling (Figure 1). By using the tailing and ssDNA ligation method (30), we optimized the sample indexing reaction to standardize the workflow of mChIP-seq (Figure S1, Supporting Information). Systematic validation confirmed high specificity, reproducibility, and stability of mChIP-seq for various sample types (Figure 1; Figure S2, Supporting Information), and demonstrated its multiplex profiling capability for 11 histone marks on 24 samples (Figure 2). Based on mChIP-seq, we further developed cf-mChIP-seq that is easy to implement for the large-scale epigenomic profiling of cf-nucleosomes (Figure 5) and employed it to analyze histone mark-associated cfDNA fragmentation patterns in 38 plasma samples from breast cancer or healthy controls, focusing on H2A.Z, H3K4me3, H3K27ac, and H3K9me3 (Figure 6).

Compared with existing sample-multiplexing methods (15–18), mChIP-seq offers at least three distinct technical advantages. First, the ssDNA ligation method used in mChIP-seq achieves higher efficiency than the dsDNA adapter ligation used in methods like Mint-ChIP and MINUTE-ChIP for DNA in crude chromatin (17, 19). In crude chromatin environments, ligation often suffers from low yields of dually-ligated fragments due to steric hindrance or limited accessibility. Consequently, these earlier methods rely on linear amplification strategies to recover sufficient single-ligated fragments for viable library construction (17, 19). Second, mChIP-seq streamlines sample indexing into a single reaction, reducing multiple steps (end-repair, A-tailing, and ligation) to just one (Table S1, Supporting Information) (30). This simplification not only reduces hands-on time and reagent costs but also minimizes sample loss and cross-contamination risks associated with multi-step processing. Finally, mChIP-seq pipelines is inherently adaptable for cell-free samples in a multiplexing way. Traditional dsDNA library preparation often fails to capture the full molecular diversity of cfDNA. In contrast, ssDNA ligation preserves single-stranded, nicked, and ultrashort cfDNA in plasma, providing a more comprehensive view of understudied cfDNA forms compared to traditional dsDNA library methods (46–48).

Analysis of mChIP-seq datasets revealed that the regulatory role of H2A.Z in gene expression is depended on its genomic deposition sites: a transcriptional activation role at promoter or enhancer regions and a transcriptional repression role at intron sites in human cells (Figure 3; Figure S9-S10, Supporting Information). Compared with non-cancerous cells, our analysis demonstrated an aberrant correlation of H2A.Z and H3K4me3 in cancer cells, which may be involved in epigenetic dysregulation of gene expression (Figure 4; Figure S11 and S12, Supporting Information). As H2A.Z overexpression has been reported across various cancer types (24–27) and is associated with poor prognosis (28, 29), we propose this overexpression may drive the abnormal deposition of H2A.Z at promoter sites. Specifically, the H2A.Z signal in genes with low H3K4me3 level was elevated in cancer cells, which was not generally observed in non-cancerous cells (Figure 4). These genes overrepresented in metabolic process, biosynthetic process, and cell cycle regulation show higher expression levels in cancer cells. It has been reported that knockdown of H2A.Z in pancreatic ductal adenocarcinoma (PDAC) cell lines could induce a senescent phenotype while overexpression could promote cellular proliferation in breast cancer cells and PDAC (25, 28). We speculate that the abnormal H2A.Z accumulation may dysregulate expression of genes essential for cancer cell proliferation, viability, and progression, however, future validation experiments involving overexpression and knock-down in cell lines will be essential to deeply investigate the impact of abnormal deposition of H2A.Z on gene expression.

Beyond epigenomic profiling of cellular nucleosomes, cell-free mChIP-seq extends high-throughput epigenomic analysis to circulating nucleosomes (Figure 5), thereby supporting the development of noninvasive liquid biopsy-based diagnostic models. The study on plasma samples with breast cancer and non-cancer controls revealed distinct histone mark-associated cfDNA fragmentation patterns among histone marks H2A.Z, H3K4me3, H3K27ac, and H3K9me3 (Figure 6; Figure S16, Supporting Information). Fragment lengths of H2A.Z-, H3K27ac-, and H3K9me3-associated cfDNA are significantly shorter in cancer samples. However, their genomic distributions are distinct, indicating their different dynamics of cf-nucleosome release and clearance. A limitation of this cfDNA study is the exclusive focus on breast cancer with small sample size. Since it has been reported that plasma DNase activity in breast cancer was elevated but reduced in colon, stomach, pancreatic, and prostate cancer (49–53), further investigation of histone mark-associated cfDNA patterns across more cancer types is warranted.

While mChIP-seq enhances efficiency, reduces cost, and lowers input requirement for epigenomic profiling of nucleosomes in cellular or cell-free forms, several considerations and limitations should be noted. As all other methods using antibodies to capture target sequences, the mChIP-seq data quality is largely dependent on the specificity of the antibody used. Increasing commercially available antibodies for histone marks with ChIP grade should improve data reliability. Another important consideration is that it is not feasible to barcode a large number of cells as typical ChIP-seq to perform chromatin immunoprecipitation, and the low input in our methods possibly result in relatively high duplicates in the data due to increasing cycles of PCR that were generally used to amplify enough DNA for sequencing. We observed relatively higher PCR duplicates in epigenomic profiles of H2A.Z, H3K4me3, H3K27ac, and H3K18ac compared with others. We inferred this is partially due to their different genomic abundance. In addition, we observed some cell lines have much less reads than others in mChIP-seq data (Table S2, Supporting Information). The difference in their barcoding efficiency may be caused by their distinct sensitivity to the same crosslink and sonication conditions used in this study.

In summary, we established mChIP-seq and its derivative, cf-mChIP-seq, as powerful tools for epigenomic profiling of histone marks with advantages of high efficiency, low cost, and low-input requirements. Our work uncovered abnormal regulatory features of H2A.Z in cancer cells and different fragmentation patterns of cfDNA associated with H2A.Z in plasma from breast cancer patients. We anticipate that mChIP-seq offers huge potential for future applications in supporting other large-scale projects like ENCODE (54), fruitENCODE (55), and RiceENCODE (56), and cf-mChIP-seq enables to expand the methodological repertoire of liquid biopsy in various clinical settings.

## Materials and Methods

### Ethics

This study for collection and usage of blood samples was approved and monitored by the Ethics Committees of the Women and Childrens Medical Center of Guangzhou Medical University . Informed consent was obtained from all individuals or their legal guardians before blood sampling. Tissue samples were handled in compliance with the guidelines (AGIS2020.04.17) provided by the Biomedical Research Ethics Committee of the Agricultural Genomics Institute at Shenzhen, Chinese Academy of Agricultural Sciences.

### Materials

Key resources used in this study were listed in Table S5.

### Cell culture

Cell lines T47D, SNB19, U251, HEK293T, and 3T3 were grown in DMEM media containing 10% Fetal Bovine Serum (FBS) and 100 U/ml Penicillin-Streptomycin (PS). Cell lines SK-MEL-2, DU145, and A498 were cultured in MEM media containing 10% FBS and 100 U/ml PS. Cell lines HCT116 and SK-OV-3 were grown in McCoy’s 5A media containing 10% FBS and 100 U/ml PS. Cell lines A549 and PC-3 were cultured in F12K media containing 10% FBS and 100 U/ml PS. Cell lines MDA-MB-468 and SW620 were grown in L15 media containing 10% FBS and 100 U/ml PS. Cell lines HL-60, K562, and RPMI-8226 were cultured in IMDM media containing 10% FBS and 100 U/ml PS. Cell lines NCI-H226, NCI-H522, CCRF-CEM, SK-MEL-28, OVCAR-8, and 786-O were grown in RPMI 1640 media containing 10% FBS and 100 U/ml PS. Cell line MOLT-4 was grown in RPMI 1640 media containing 1.5 g/L NaHCO_3_, 10% FBS, and 100 U/ml PS. Cell lines NCI-H23 and NCI-H460 were grown in RPMI 1640 media containing 10% FBS, 2 mM L-Glutamine, and 1 mM Sodium Pyruvate. All cells were grown at 37°C with 5% CO_2_ in a humidified atmosphere. Yeast strain BY4741 was grown in YPD media (1% yeast Extract, 2% Peptone and 2% glucose) at 28°C with 200 rpm on a shaker.

### Sample fixation

For attached cell lines, cells were grown on 10-cm plates. When confluence reached to 80%-90%, culture media were aspirated and cells were fixed with 10 ml of freshly prepared cross-linking buffer (50 mM HEPES-KOH PH 8.0, 1 mM EDTA, 1 mM EGTA, and 150 mM NaCl, and 1% formaldehyde) at room temperature for 10 min with moderate shaking. Immediately after, glycine with 0.125 M final concentration were added to stop the fixation at room temperature for 5 min. Then, cells were washed with ice-cold PBS twice, scraped from the plates in 10 ml of ice-cold PBS containing 5 mM EDTA and 1×protease inhibitors, and transferred to a 15 ml tube on ice. Cells were pelleted by centrifugation at 500× g, 4°C for 5 min. The pellet was transferred to a 1.5 ml tube with 1 ml of ice-cold PBS containing 5 mM EDTA and 1×protease inhibitors and cells were collected by centrifugation at 500× g, 4°C for 5 min. After the supernatant removing, fixed cell pellets were snap-frozen by liquid nitrogen and stored at −80°C before use.

For suspension cell lines, cells were grown in T75 flasks. When cell density reached to ∼1×10^6^ cell/ml, cells were transferred to a 15 ml and collected by centrifugation at 500× g, 4°C for 5 min. The pellet was resuspended in 500 μl of PBS and subsequently fixed with 10 ml of freshly prepared cross-linking buffer at room temperature for 10 min while gently inverting in a rotator. Immediately after, glycine with 0.125 M final concentration were added to stop the reaction at room temperature for 5 min. Then, cells were collected by centrifugation at 500× g, 4°C for 5 min, resuspended in 1 ml of ice-cold PBS containing 5 mM EDTA and 1×protease inhibitors, transferred to a 1.5 ml tube, and washed with 1 ml of ice-cold PBS containing 5 mM EDTA and 1×protease inhibitors twice. The fixed cells were snap-frozen by liquid nitrogen and stored at −80°C before use.

For frozen tissues stored at −80°C, the samples were ground to a fine powder in liquid nitrogen and fixed with 10 ml of freshly prepared cross-linking buffer for 10 min at room temperature. Immediately after, glycine with 0.125 M final concentration were added to quench the fixation at room temperature for 5 min. Then, fixed tissues were collected by centrifugation at 500× g, 4°C for 5 min and washed with 1 ml of ice-cold PBS containing 5 mM EDTA and 1×protease inhibitors twice. The fixed tissues were snap-frozen by liquid nitrogen and stored at −80°C before use.

For yeast, cells were grown at 28°C with 200 rpm on a shaker overnight. To fix cells, formaldehyde was directly added to the culture with a final concentration of 1% and cells were cross-linked at 25°C for 20 min with 200 rpm on a shaker. Glycine with 0.125 M final concentration were added to quench the reaction at 25°C for 5 min. Then, cells were collected by centrifugation at 500× g, 4°C for 5 min and washed with ice-cold PBS containing 5 mM EDTA and 1×protease inhibitors three times. The fixed cells were snap-frozen by liquid nitrogen and stored at −80°C before use.

### Fragmentation and quantification

For mammalian cell, 2-5 million cell aliquots were thawed on ice and resuspended in 1 ml of cold Cell lysis buffer (CLB, 50 mM Tris-HCl H 8.0, 150 mM NaCl, 1mM EDTA, 1 mM EGTA, 10% Glycerol, 0.5% IGEPAL®CA-630, 0.25% Triton X-100, and 10 mM sodium butyrate) with 1× protease inhibitors for 20 min on ice with inversion every 4 min. The nuclei were pelleted by centrifugation at 2,000× g for 5 min at 4°C. The supernatant was removed and the pellet was resuspended in 130 μl of Nuclear lysis buffer (10 mM Tris-HCl pH 8.0, 1 mM EDTA, 1 mM EGTA, and 0.4% SDS; 1× protease inhibitors were added before use). Then, nuclei were sonicated using Covaris SE220 (75 W, 15% Duty factor, 1,000 cycles per burst) for 5 min at 4°C to obtain fragments. Then, the solution was centrifuged at 16,000× g for 10 min at 4°C to pellet the insoluble cell debris. For each sample, 20 μl of chromatin extracts were collected to check sonication results and quantifying DNA concentration in the extracts, and the rest were stored in single-use aliquots (about 0.1 million cells per aliquot) at −80°C.

For tissue, about 50 mg of samples were thawed on ice and resuspended in 1 ml of cold CLB with 1× protease inhibitors and homogenized using a Dounce homogenizer on ice. The tissue homogenate was then pelleted by centrifugation at 2,000× g for 5 min at 4°C. After removing the supernatant, the pellet was resuspended in 130 μl of Nuclear lysis buffer and fragmented as mammalian cell.

For yeast, chromatin was extracted according to prior protocols with some modifications (57, 58). In brief, frozen cells overnight cultured in 50 ml media were thawed on ice for 30 min and resuspended in 1 ml of cold yeast ChIP lysis buffer (50 mM Tris-HCl pH 7.5, 1 mM EDTA, 1% IGEPAL®CA-630) with 1 mM phenymethylsulfonylfloride (PMSF) and 1× protease inhibitors. Then, the suspension was transferred to a 2 ml tube with 500 μl of cold glass beads and lysed by bead beating in a Cryogenic Grinder for 6 cycles with 30 s on / 60 s off at 4°C. The lysate was transferred to a 1.5 ml tube and centrifuged at 16,000× g for 20 min at 4°C. The supernatant was discarded and the pellet was resuspended in 130 μl of ChIP binding buffer (CBB, 10 mM Tris-HCl pH 7.5, 150 mM NaCl, 1 mM EDTA, and 1% Triton X-100, 0.1% SDS) with 1× protease inhibitors. Then, the lysate was sonicated using Covaris SE220 (75 W, 20% Duty factor, 1,000 cycles per burst) for 15 min at 4°C to obtain DNA fragments between 100-500 bp in size. After sonication, cell debris was pelleted by centrifugation at 16,000× g for 10 min at 4°C. and the chromatin extracts in the supernatant were stored in single-use aliquots (about cells in 5 ml media per aliquot) at −80°C after taking out 20 μl for DNA quantification.

To quantify DNA concentration, 20 μl of chromatin extracts were mixed with 124 μl of Elution buffer (50 mM Tris-HCl pH 8.0, 10 mM EDTA, and 0.5% SDS) and incubated at 65°C for 3 hours to reverse-cross-link the protein-DNA complexes. Subsequently, the eluate was digested with 10 μg of RNase A at 37°C for 1 hour to remove RNA and then 100 μg of proteinase K at 65°C for 2 hours to remove proteins. DNA concentration was quantified by Qubit™ 1× dsDNA High Sensitivity (HS) Kit, purified by phenol:chloroform:isoamyl alcohol extraction, precipitated with ethanol, and resuspended in TE buffer for fragment size check by gel electrophoresis.

### mChIP-seq for cell and tissue samples

Chromatin extracts were taken out from -80°C, thawed on ice, and centrifuged at 16,000× g, 4°C for 15 min. The supernatant was then transferred to a clean tube, diluted by 360 μl of ChIP Dilution buffer (10 mM Tris-HCl pH 8.0, 1 mM EDTA, and 1 mM EGTA; 1× protease inhibitors were added before use), added Triton X-100 with a final concentration of 1%, and incubated at 37°C for 20 min to quench the SDS in the solution. To index chromatin DNA 3’ ends, the fragmented chromatin solution contained about 200 ng DNA quantified by Qubit™ as above mentioned were mixed with 4 μl of P7TruSeqHT/P7SPlintC (Phosphate-CAGCGATCGACNN NNAGATCGGAAGAGCACACGTCTGAACTCCAGTCA-ddC/ GTCGATCGCTGCCCCCC-NH2 C6, NNNN: barcodes with four different deoxynucleotide;), 10 μl of 10× T4 DNA ligase buffer, 2.5 μl of 10 mM dGTP, 10 Weiss units of T4 DNA ligase, 20 units of T4 Polynucleotide Kinase (T4 PNK), 40 units of Terminal Deoxynucleotidyl Transferase (20 U/μl) (TdT), and ddH_2_O to a total volume of 100 μl. For Pol II profiling in this study, the reaction volume was scaled up to 200 μl with 400 ng DNA of fragmented chromatin solution. Then, the mixture was incubated at 37°C for 1 hour, and the reaction was stopped by adding 3 μl of 0.5 M EDTA and 2 μl of 10% SDS, followed by incubated at 37°C for 20 min.

Barcoded samples were pooled, one tenth volume of the mixed solution was stored at -20°C for input library preparation, nine tenth volume of the mixed solution were diluted by adding equal volume of 2× CBB (20 mM Tris-HCl pH 7.5, 300 mM NaCl, 2 mM EDTA, and 2% Triton X-100). Considering the genome size of budding yeast is much smaller than the size of human and mouse, barcoded fragmented chromatin of extracts contained about 20 ng DNA of BY4741 were taken for mChIP-seq experiment of which samples from human cell lines HEK293T and HeLa, mouse cell line 3T3, and budding yeast strain BY4741 were pooled (Figure 1). Before splitting, the chromatin pool was pre-cleared by adding the washed Pierce™ ChIP-grade Protein A/G Magnetic Beads and incubated at 4°C in a rotator by rotating at 10 rpm for at least 2 hours. Pierce™ ChIP-grade Protein A/G Magnetic Beads were washed with 1 ml of 1× CBB twice by gently inverting in a rotator for 3∼5 min per time at 4°C. The volume of beads was calculated according to the number of samples. In this study, we used 5 μl of beads per sample to pre-clear the chromatin pool. Subsequently, beads were removed by using a magnetic rack

and the supernatant was split for immunoprecipitation. The amount of indexed chromatin we put for each antibody included about 400 ng DNA according to the DNA quantification as above described in this study. To perform immunoprecipitation, 2 μg of antibody was added to each tube. Then, Chromatin and antibody in 500 μl volume of 1×CBB were rotated overnight at 4°C. The next day, 10 μl of pre-washed Protein A/G Magnetic beads were added to each immunoprecipitation tube, incubated for 3 hours at 4°C to capture the immuno-complexes by gently rotating, and washed twice with each of the following buffers: 500 μl of low-salt washing buffer (0.1% SDS, 1% Triton X-100, 2 mM EDTA, 150 mM NaCl, 20 mM Tris-HCl, pH 7.5), 500 μl of high-salt washing buffer (20 mM Tris-HCl, pH 7.5, 0.1% SDS, 1% Triton X-100, 2 mM EDTA, 500 mM NaCl), 500 μl of LiCl washing buffer (250 mM LiCl, 1% NP-40, 1% Na deoxycholate, 1 mM EDTA, 10 mM Tris-HCl, pH 7.5), and 500 μl of TE buffer (10 mM Tris-HCl, pH 8.0, 1 mM EDTA).

To elute DNA, we added 40 μl of Elution buffer (10 mM Tris-HCl pH 8.0, 1 mM EDTA, 0.5% SDS, and 150 mM NaCl) to the beads followed by incubation at 37°C for 15 min. Subsequently, beads were removed by using a magnetic rack and the supernatant was transferred to a new tube. To revert formaldehyde crosslinking, we added proteinase K with a final concentration of 1 mg/ml to the supernatant and incubated it at 65°C for at least 2 hours. Then, 1.8× volume of SPRIselect beads according to the manufacturer’s instruction or a standard phenol-chloroform extraction procedure were used to purify ChIPed DNA.

To prepare sequencing library, the purified ChIPed DNA was resuspended in 18 µl of water and extended in a 40 µl reaction containing 1× KAPA HiFi HotStart ReadyMix and 0.25 µM P7TruSeqHTRev (GTGACTGGAGTTCAGACGTGTGCTCTTCCGATC) at 98 for 30 sec and 63°C for 5 min followed by purification with 1.2× volume of SPRIselect™ beads. Then, the purified DNA was ligated to the second adapter in a 40 µl reaction (1× T4 DNA ligase buffer, 0.5 µM P5TruSeqHT/P5SPlint mixture (ACACTCTTTCCCTACACGACGCTCTTCCGATCT/NH2C6-AGATCGGAAGAGC), and 10 Weiss units of T4 DNA ligase) at 25°C for 30 minutes and purified with 1.0× volume of SPRIselect™ beads. Lastly, library was amplified in 50 μl of PCR reaction Mix containing eluted DNA, 0.5 μM of P5 index primer, 0.5 μM P7 index primers, and 25 μl of KAPA HiFi HotStart ReadyMix (2×) for 10-17 cycles (each cycle with 20 s at 98°C, 30 s at 58°C, 1 min at 68°C). DNA of 200-700 bp was isolated using a double SPRIselect bead cleanup (0.6× and 0.85× volume). Libraries were quantified by the Qubit™ 1× dsDNA High Sensitivity (HS) Kit, qualified by Fragment Analyzer (Advanced Analytical), and subjected to 150-bp-paired-end sequencing (30).

### cf-mChIP-seq for blood samples

Blood samples were collected in K3 EDTA tubes, transferred immediately to ice, and 1× protease inhibitor cocktail and 10 mM EDTA were added as reported in (5, 7). Plasma was isolated from blood cells whin 4 hours after sample collection. The whole blood was centrifuged at 1500 × g, 4°C for 10 min. The supernatant was transferred to a new tube and centrifuged at 3000 × g, 4°C for 10 min. The plasma extraction was then aliquoted, flash frozen and stored at -80°C before use.

To perform sample indexing, 100 μl of plasma were mixed with 8 μl of P7TruSeqHT/P7SPlintC, 20 μl of 10 × T4 DNA ligase buffer, 2.5 μl of 10 mM dGTP, 10 Weiss units of T4 DNA ligase, 20 units of T4 Polynucleotide Kinase (T4 PNK), 40 units of TdT (20 U/μl) and ddH2O to a total volume of 200 μl (30). Then, the reaction was incubated at 37°C for 1 hour and stopped by adding 3 μl of 0.5 M EDTA and 2 μl of 10% SDS and incubating at 37°C for 20 min. During the sample indexing, the antibody-bead complexes were prepared by incubating 2 μg of each antibody with pre-washed 20 μl of Pierce™ ChIP-grade Protein A/G Magnetic Beads at 4°C for at least 3 h before use in a rotator.

After sample indexing, barcoded samples were pooled together and centrifuged at 16,000 × g, 4°C for 10 min. The supernatant was transferred to a new tube and pre-cleared with 50 μl of Pierce™ ChIP-grade Protein A/G Magnetic Beads at 4°C for 2 h in a rotator. Subsequently, beads were removed by using a magnetic rack and the supernatant was split for ChIP. To perform ChIP, the pooled samples and antibody-bead complexes were mixed and rotated overnight at 4°C. The next day, the chromatin-antibody-bead complexes were washed with the buffers twice for each in the following order: 500 μl of low-salt washing buffer (0.1% SDS, 1% Triton X-100, 2 mM EDTA, 150 mM NaCl, 20 mM Tris-HCl, pH 7.5), 500 μl of high-salt washing buffer (0.1% SDS, 1% Triton X-100, 2 mM EDTA, 500 mM NaCl, 20 mM Tris-HCl, pH 7.5), 500 μl of LiCl washing buffer (250 mM LiCl, 1% IGEPAL®CA-630, 1% Na deoxycholate, 1 mM EDTA, 10 mM Tris-HCl, pH 7.5), and 500 μl of TE buffer (10 mM Tris-HCl, pH 8.0, 1 mM EDTA).

After wash, beads were resuspended in 40 μl of Elution buffer (10 mM Tris-HCl pH 8.0, 1 mM EDTA, 0.5% SDS, and 150 mM NaCl) with 1 mg/ml proteinase K and incubated for 1 h at 55°C to elute ChIPed DNA. Then, 1.8 × volume of SPRIselect beads according to manufacturer’s instructions were used to purify ChIPed DNA. Then sequencing libraries was prepared as above mentioned.

### Data processing and analysis for mChIP-seq

Sequencing reads from mChIP-seq were demultiplexed to samples by the barcodes at the 5’ end of Read2 using fastq-multx (version 1.4.2) (https://github.com/brwnj/fastq-multx) with a mismatch setting of 0. Then, reads of each sample with adapter contamination and low quality were trimmed and filtered using fastp with parameters as -f 10 -t 5 -F 30 -T 5 --complexity_threshold=30 –n_base_limit=5 --length_required 60 (version 0.23.2) (59) and aligned to the human reference genome hg38, mouse reference genome mm10, pig reference genome Sscrofa11 or budding yeast genome sacCer3 using bowtie2 (version 2.2.5) with default parameters (60). Uniquely mapped reads with mapping quality no less than 30 were kept using samtools (version 1.9) (60) and duplicates were removed using picard MarkDuplicates (v2.26.10, http://broadinstitute.github.io/picard). External downloaded datasets were directly used to map to reference genome using bowtie2 (version 2.2.5) with default parameters and then processed identically to own generated datasets (Table S3).

The processed data sets were used to obtain peak regions using MACS2 (v2.2.6) (61) with broad option -b and a cutoff of q=0.01 for yeast samples. For mammalian samples, BAM files of replicates were merged and the merged BAM files were used to define enrichment regions using SICER2 software (62) with a window size setting of 200 bps, a gap setting of 400 bps, a FDR setting of 0.001 for “sharp” marks (H2A.Z, H3K4me1, H3H4me2, H3K4me3, H3K27ac, and H3K18ac), and a window size setting of 200 bps, a gap setting of 600 bps, an FDR setting of 0.001 for “broad” marks (H3K36me3, H3K27me3, and H3K9me3) to Input data (fold-change > 2).

To obtain enrichment sites of H2A.Z for each cell line (Figure 4a), SICER2 was used to call enrichment regions of the individual replicates with a window size setting of 200 bps, a gap setting of 400 bps, an FDR setting of 0.01 to Input data (fold-change > 2). Then, enrichment regions of the individual replicates were concatenated, sorted using the BEDtools (v2.30.0,73) SortBed function. Common intervals among replicates were identified by the BEDtools Multiinter function and selected by awk command. These common intervals with bookended intervals were combined into a single by the BEDtools Merge function to obtain the final enrichment sites in BED file format of each sample for subsequent analysis. To get the total enrichment sites of H2A.Z for the 24 cancer cell lines, the final enrichment sites in BED files of these cell lines were concatenated, sorted using the BEDtools (v2.30.0,73) SortBed function, and overlapping regions (by at least 1 bp) were merged by the BEDtools Merge function. Public data sets were processed identically to own generated datasets, except that sites with fold-change not less 3 were retained as enrichment sites for samples without replicates. To annotate the genomic distribution of H2A.Z enrichment sites, R package ChIPseeker (version 1.26.2) (63) were used with tssRegion=c(-1000,1000) and the priority order (promoter > exon > intron > intergenic) when a single peak spanned more than two genomic features.

### Data processing and analysis for cf-mChIP-seq

Sequencing reads from cf-mChIP-seq were demultiplexed to samples as mChIP-seq described above. Then, reads of each sample with adapter contamination and low quality were trimmed and filtered using fastp with parameters as --complexity_threshold=30 –n_base_limit=5 --length_required 60 (version 0.23.2) (59) and aligned to the human reference genome hg38, using bowtie2 (version 2.2.5) with default parameters (60). Uniquely mapped reads with mapping quality no less than 30 were kept using samtools (version 1.9) (60) and duplicates were removed using picard MarkDuplicates (v2.26.10, http://broadinstitute.github.io/picard).

For peak calling, the processed bam files of cancer samples and non-cancer controls were merged with samtools, respectively. Then merged files were used to obtain peak regions using MACS2 with --nomodel -q 0.01 --keep-dup all --fe-cutoff 5 for H2A.Z, H3K4me3, and H3K27ac and with –nomodel -b --broad-cutoff 0.05 --fe-cutoff 2 for H3K9me3.

Fragment size frequency and genome-wide fragmentation pattern analysis were performed as described previously (64, 65). In brief, the total number of fragments for each sample were used to normalize fragment size frequency. Fragments with size in 90-250 bp and 251-450 bp were assigned as mono-nucleosome fractions and di-nucleosome fractions, respectively. The median frequency for each fragment size was calculated in each group. For fragmentation pattern analysis at the genome-wide level, fragments were assigned to 5-Mb adjacent, non-overlapping bins using the ‘bedtools intersect’ tool. Bins with an average GC content <0.3 and an average mappability <0.9 were excluded as previously described(65, 66). Differently, we focused on the fragment ratios of the number of mono-nucleosome to di-nucleosome for each 5-Mb bin in this study.

### Data processing and analysis for RNA-seq

For RNA-seq data, reads were aligned to hg38 using HISAT2 (version 2.2.1) with default parameters (67). Then, raw counts for the feature of genes were extracted by featureCounts with the -B parameter to summarize fragments that have both ends successfully aligned (version 2.0.3) (68). R package edgeR was used to normalize the data with TMM method, and RPKM was calculated by the rpkm function in edgeR (69).

### Correlation analysis

To perform correlation analysis of signals in peaks, raw reads of peaks merged from all samples were extracted and normalized to count per million (CPM). For correlation analysis for signals in genome-wide, the processed BAM files were converted to normalized coverage files (bigWig) with 5-bp bins and without blacklisted regions (https://github.com/Boyle-Lab/Blacklist/) using bamCoverage from deepTools (version 3.5.1) (70) and deepTools plotCorrelation function was used to compute correlation coefficients depicting as a clustered heatmap. For correlation analysis shown in Figure 6d and 6e, we selected samples with usable reads around 0.1 Mio or more based on the H2A.Z dataset (not less than 0.08 Mio to include samples as more as possible), and finally retained 25 samples (15 patients and 10 healthy controls). Then, we downsampled the number of usable reads to 0.1 Mio for samples with reads more than 0.1 Mio. H2A.Z peaks in promoter overlapping among mammary epithelial cell, and breast cancer cell lines MDA-MB-468 and T47D (n = 4,395) were used for Spearman correlation analysis.

### GO enrichment analysis

GO enrichment analysis was performed using DAVID (The Database for Annotation, Visualization and Integrated Discovery) (71). Terms of GOTERM_BP_2 were used in this study.

### Data visualization

Snapshots of the data sets were constructed by the Integrative Genomics Viewer (IGV) (72).

### Data Availability

Sequencing data have been deposited at Gene Expression Omnibus (GEO) (accession number GSE280574).

### Statistical Analysis

Statistical tests for all data were conducted using R (version 4.2.2). Unless otherwise stated, all data are presented as mean ± SD. Comparisons between two groups were performed using a Paired Samples t-Test, Welch’s t-test, or Mann-Whitney U tests mentioned in legends. Statistical significance was defined as p < 0.01 or p < 0.001.

## Acknowledgements

We greatly appreciate all The Wei Xu Lab members for useful discussions. We would like to express our deep appreciation to Zhonglin Tang Lab for providing pig tissues. This work was supported by National Key R&D Program of China (Grant No. 2021YFF1000600), Basic Research Center, Innovation Program of Chinese Academy of Agricultural Sciences (CAAS-BRC-AFIS-2025-02), Central Public-interest Scientific Institution Basal Research Fund (No. Y2022QC33), and National Natural Science Foundation of China (Grant No. 32071437 and 31900302).

## Authors contributions

Xu Wei and Jie Li conceived the study and supervised the experiments and data analysis. Changbin Sun performed mChIP-seq experiments and data analysis. Qinkai Zhang, Jianli Yan, Xilin Wang, Chunxiao Zhang and Yujie Li prepared materials. Changbin Sun wrote the manuscript. Wei Xu reviewed and modified the manuscript. All authors read and approved the final manuscript.

## Competing interests

The authors declare no competing interests.

## Supplementary Information

### Supplementary Figure Legends

**Figure S1.**
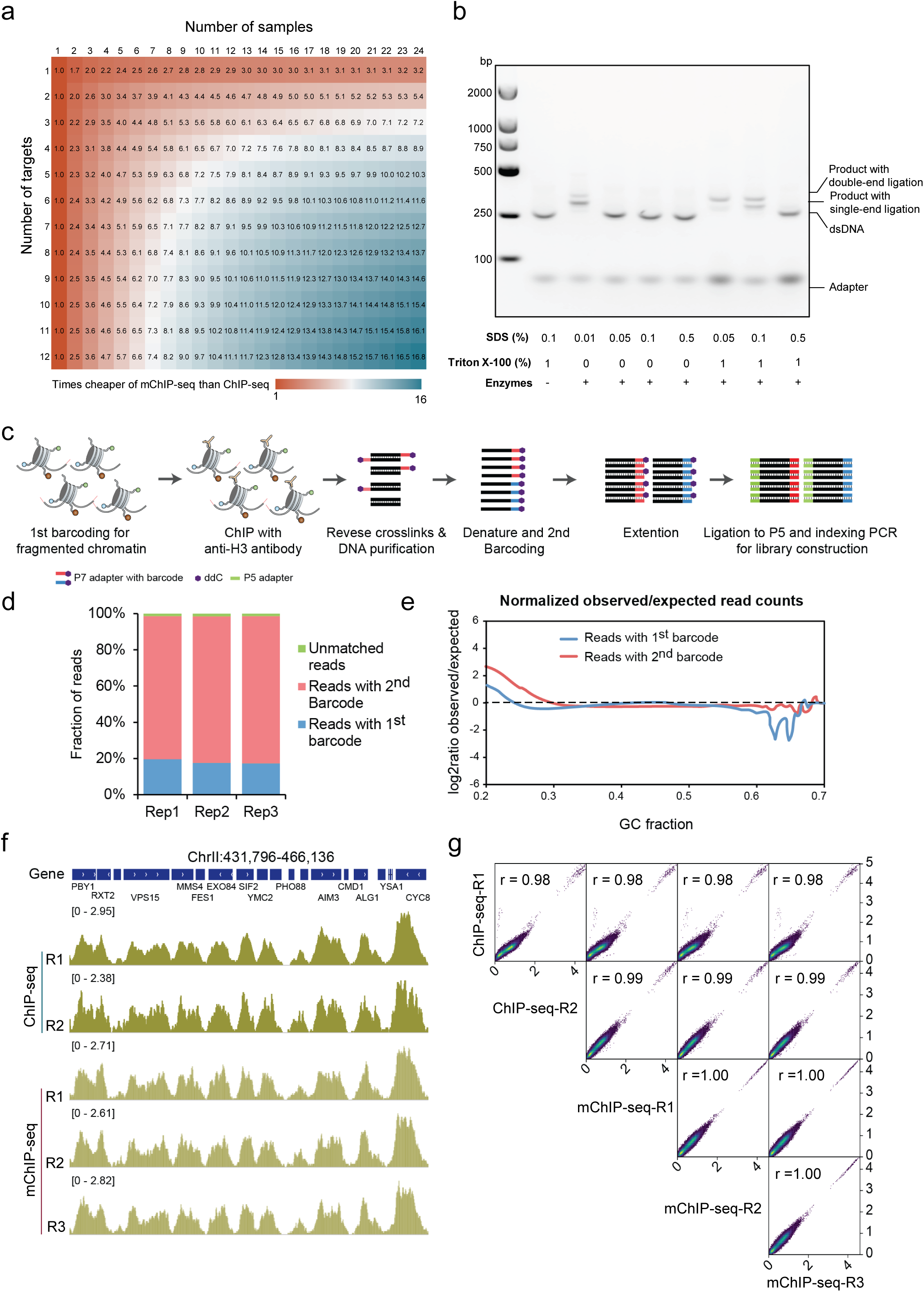
Optimization of the tailing and ssDNA ligation method for sample barcoding. a) Heatmap showing cost reduction of mChIP-seq for multiplex and multifactorial epigenomic profiling. The times cheaper of mChIP-seq than ChIP-seq was calculated by dividing cost of ChIP-seq by the cost of mChIP-seq for the same number of samples and targets. b) Gel image showing influences of SDS and Triton X-100 on the ligation efficiency. c) Schematic diagram showing workflow for ligation efficiency assessment. d) Bar plot showing barcoding efficiency for BY4741 chromatin. e) GC-bias plots showing bias of read distribution across different GC content regions. fc) Genome browser tracks comparing mChIP-seq and ChIP-seq for profiling H3K4me1 in budding yeast. g) Pairwise heatscatter plots showing correlations of genomic H3K4me1 signals in budding yeast profiled by mChIP-seq and ChIP-seq. Size of the bins:100 bp.

**Figure S2.**
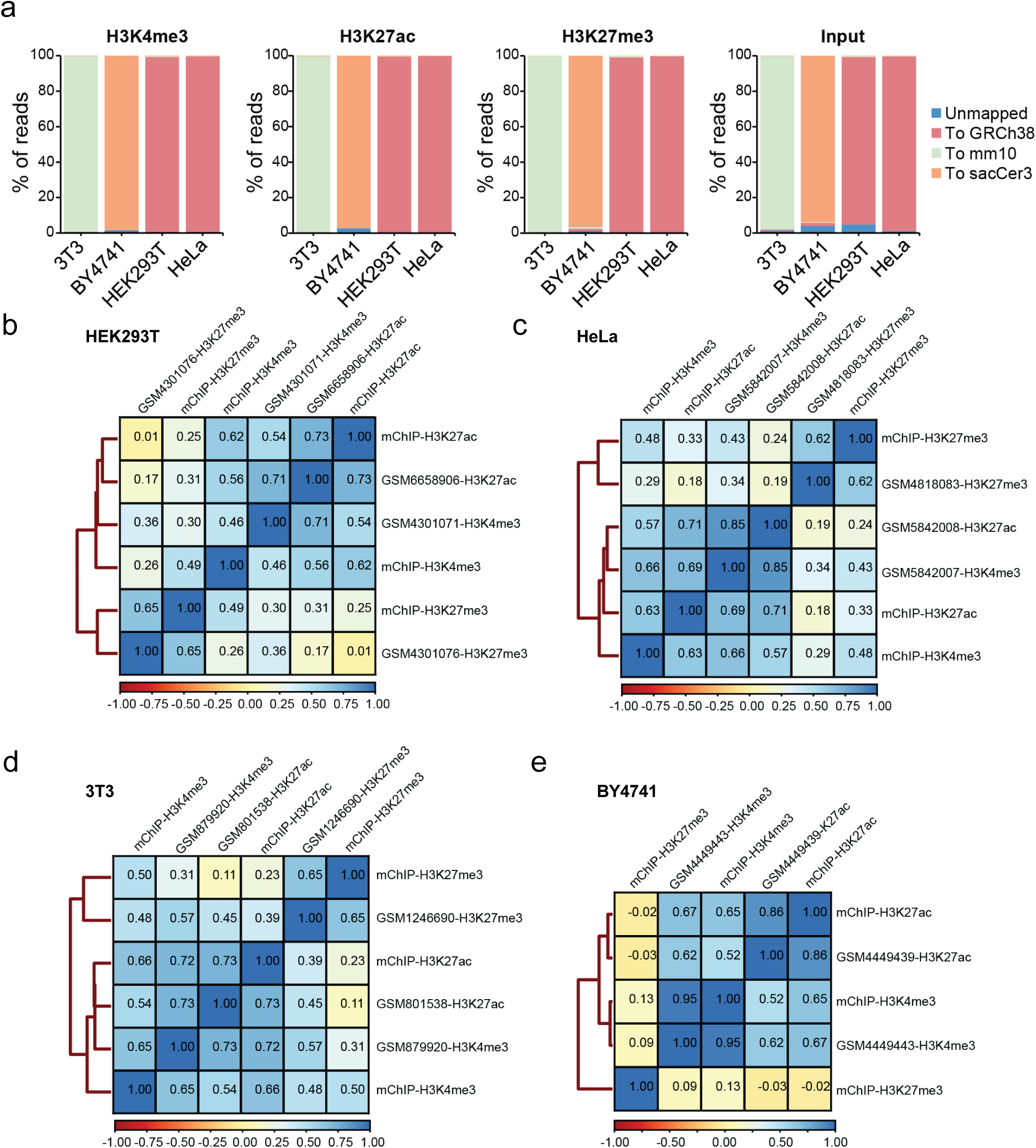
Validation of mChIP-seq for cell samples. a) Bar plots depicting percentage of reads mapping to human genome (hg38), mouse genome (mm10), and yeast genome (sacCer3) in mChIP-seq assays for profiling histone modifications of H3K4me3, H3K27ac, and H3K27me3 in human cell lines HEK293T and HeLa, mouse cell line 3T3, and budding yeast strain BY4741. Samples were indexed, pooled and split for profiling each modification. The data are shown as mean ± SD of four replicates. b) to e) Heatmap comparing genomic signal correlations between mChIP-seq and ChIP-seq for profiling H3K4me3, H3K27ac, and H3K27me3 in HEK293T (b), HeLa (c), 3T3(d), and BY4741 (e). Average signals of the replicates in mChIP-seq were used. The correlation coefficients were computed by the Spearman method. Bin size was set to 10 kb for HEK293T, HeLa, and 3T3, and 1000 bp for BY4741.

**Figure S3.**
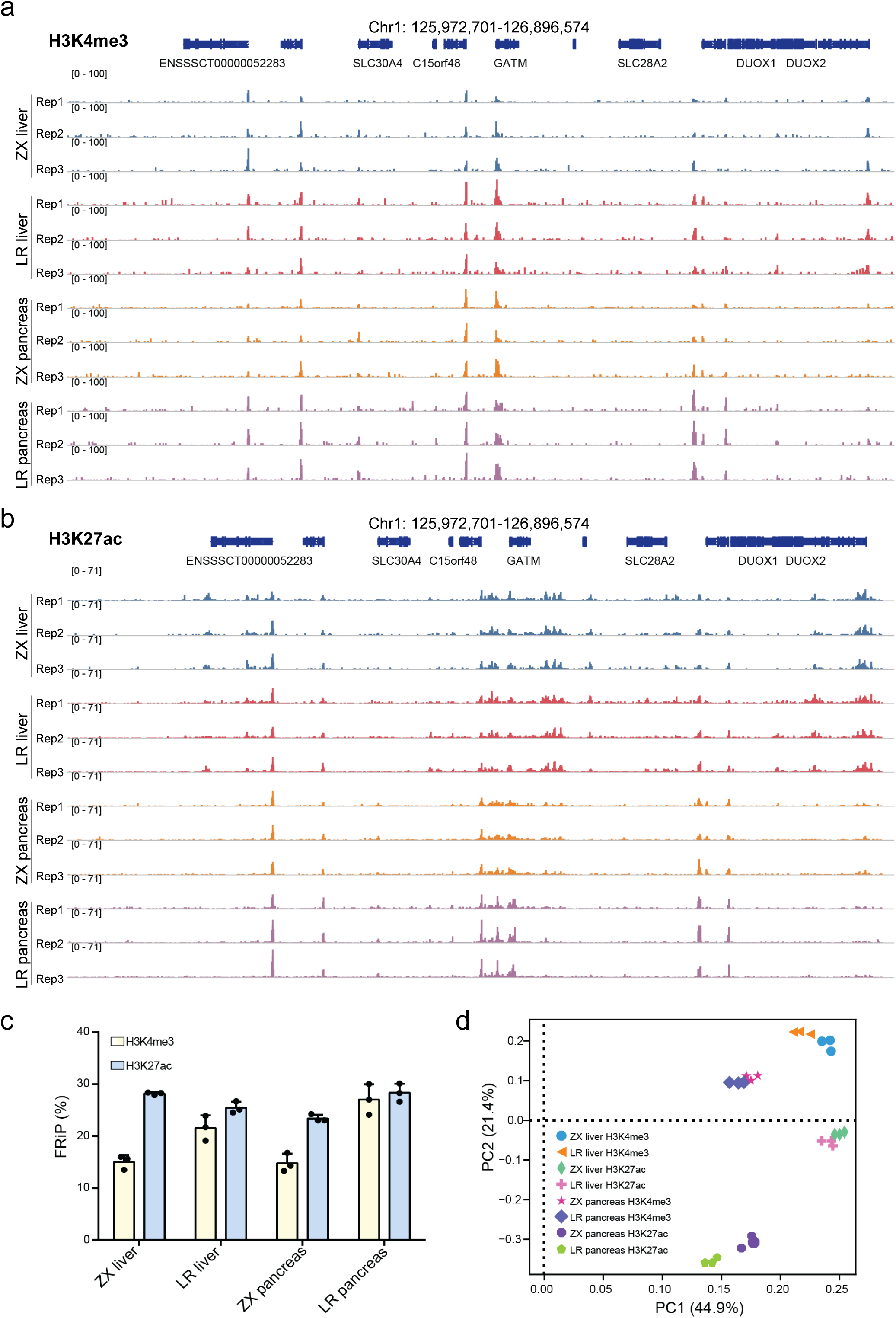
Validation of mChIP-seq for tissue samples. a) and b) Representative genome browser tracks of H3K4me3 (***a***) and H3K27ac (b) for liver and pancreas tissues from two pig breeds. c) Bar plot showing FRiP of H3K4me3 and H3K27ac for pig tissues. Merged peaks of each modification were used for calculating FRiPs. d) Principal component analysis (PCA) depicting variations of samples. Samples of liver and pancreas with three replicates from two pig breeds were pooled and divided for H3K4me3 and H3K27ac modifications. ZX: Zangxiang breed; LR: Landrace breed.

**Figure S4.**
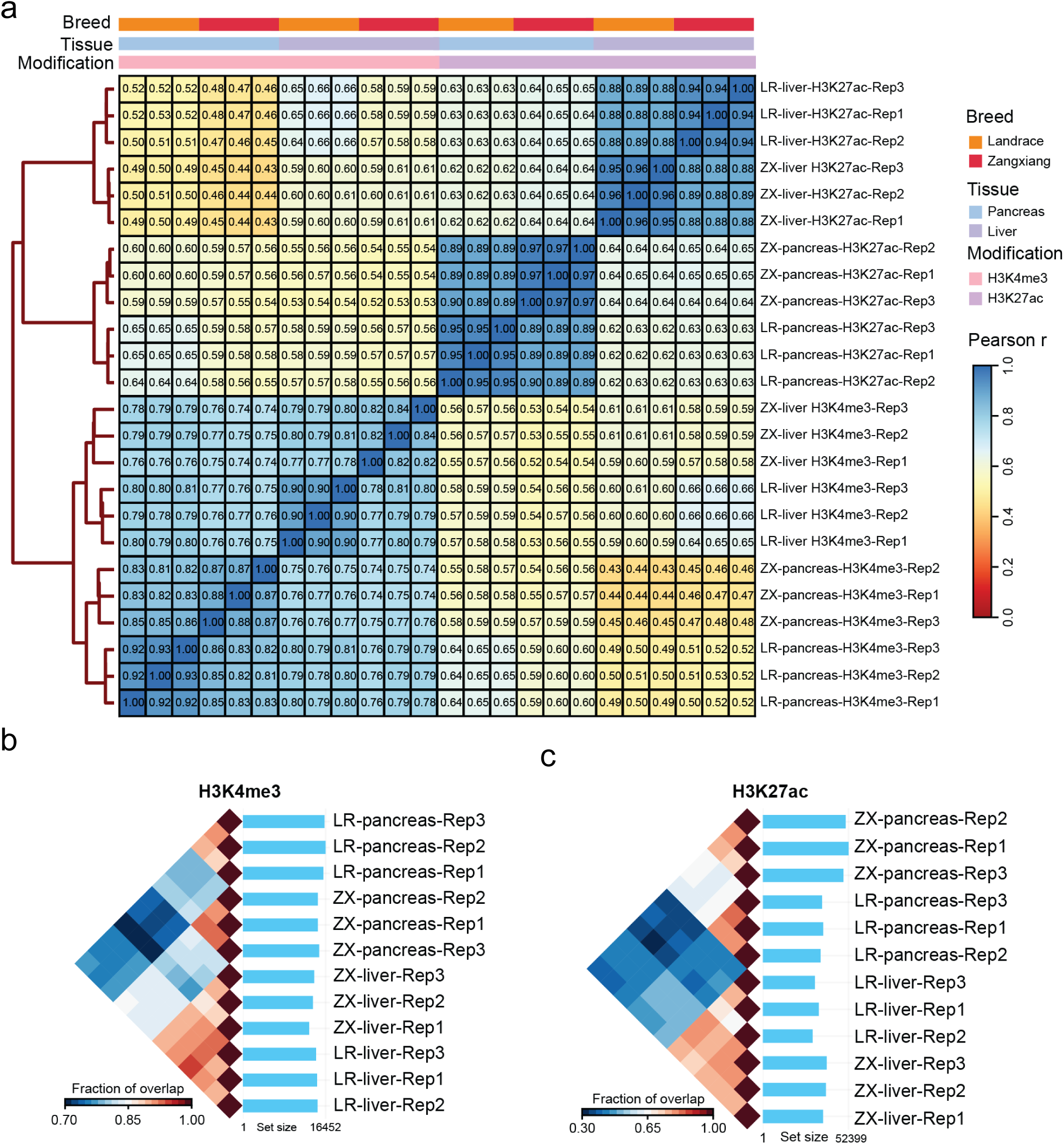
High reproducibility of mChIP-seq. a) Heatmap showing signal correlations among replicates at genome-wide. Bin size: 10 kb. b) and c) Pairwise intersection of enrichment sites for H3K4me3 (b) and H3K27ac (c).

**Figure S5.**
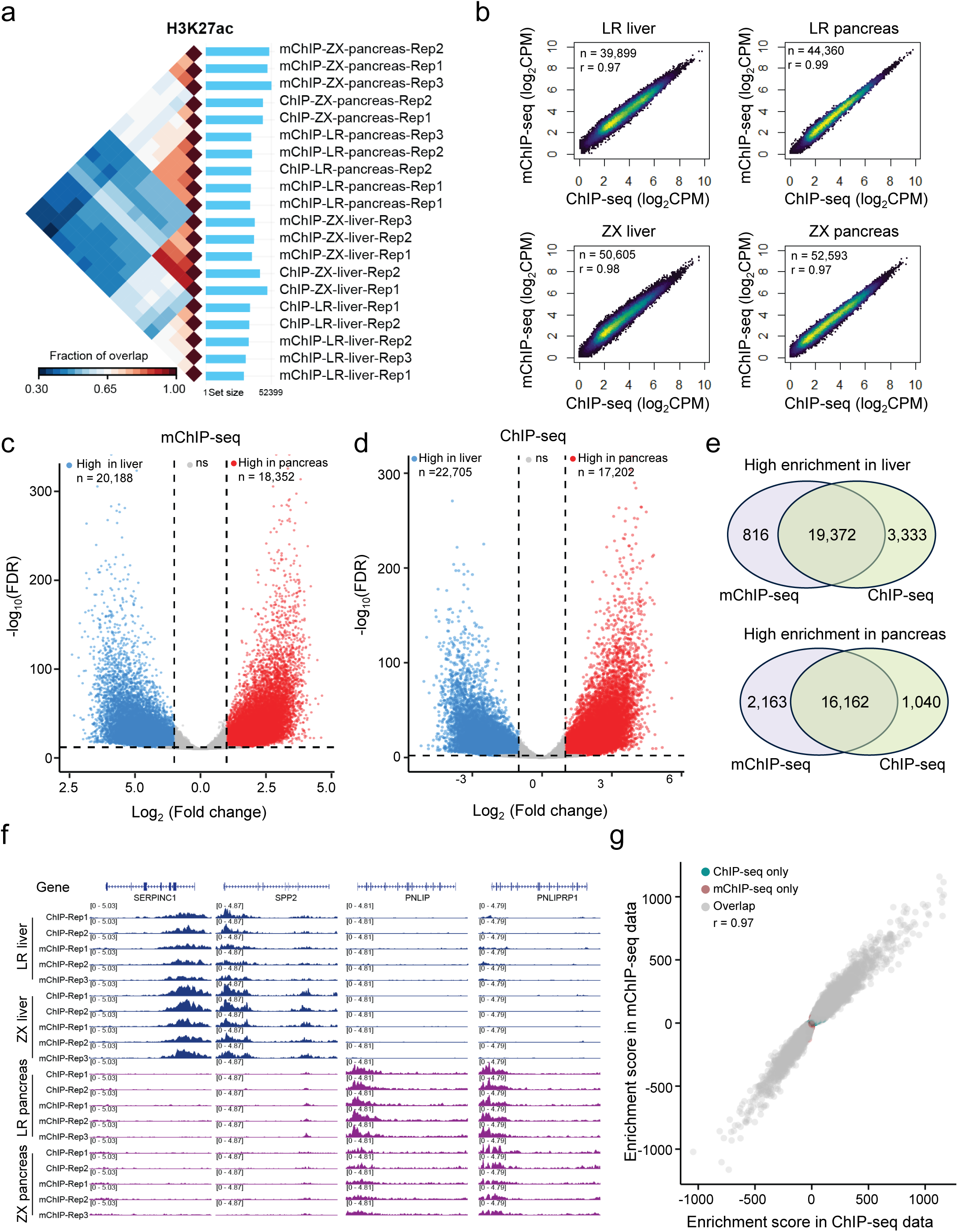
Performance of mChIP-seq for comparative analysis. a) Pairwise intersection showing fractions of overlapping peaks. ZX: Zangxiang breed; LR: Landrace breed. b) Heatscatter plots showing the correlations of peaks overlapping between mChIP-seq and ChIP-seq. n: number of peaks; r: Pearson correlation coefficient. c) and d) Volcano plot showing DESs of H3K27ac between the liver and pancreas tissues identified by mChIP-seq (c) and ChIP-seq (d) datasets. The number of DESs is given. DESs: FDR ≤ 0.01 and absolute value of log2(fold change) ≥ 1. e) Venn diagrams showing overlapping DESs shared between mChIP-seq and ChIP-seq datasets. f) Representative tracks of H3K27ac signals around tissue-specific genes. g) Scatter plot showing the correlation of enrichment score between mChIP-seq and ChIP-seq datasets in DESs. Enrichment score: fold chang multiply -log10(FDR); r: Pearson correlation coefficient.

**Figure. S6.**
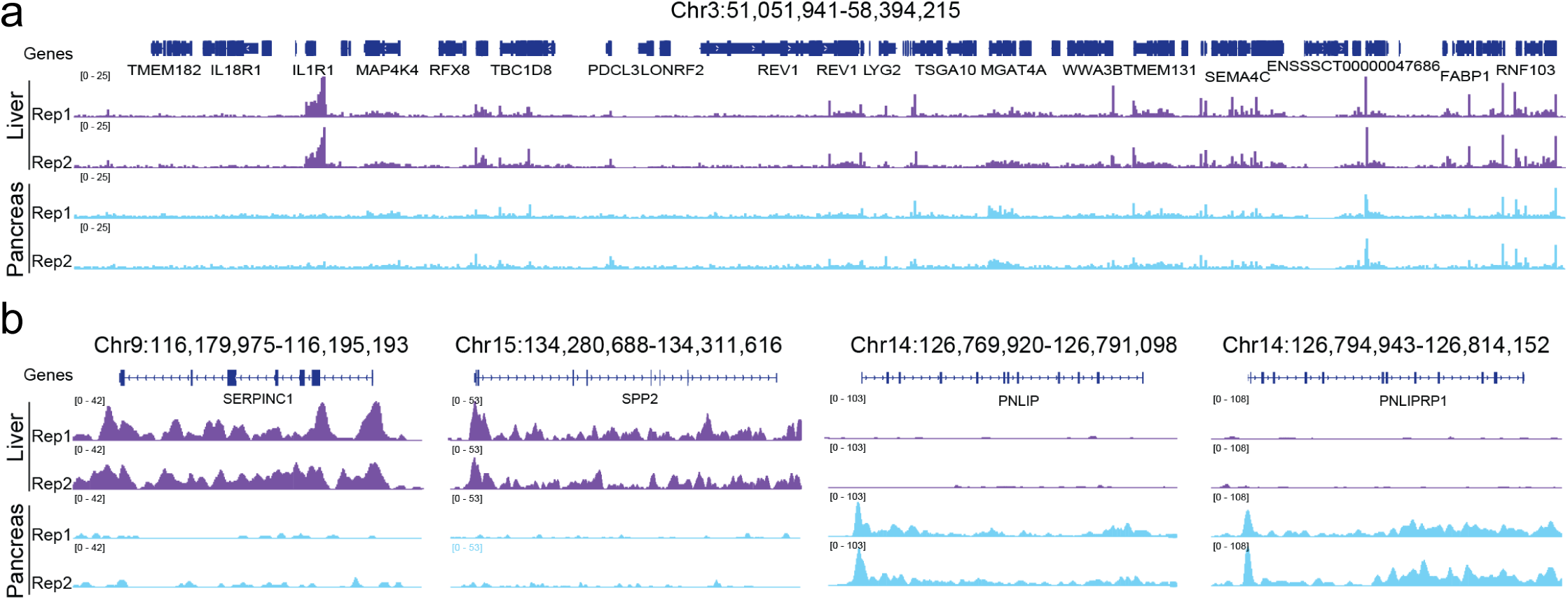
mChIP-seq for RNA Pol II profiling. a) Genomic track of RNA Pol II signal in a large genomic region. b) Genomic tracks of RNA Pol II signals around representative tissue-specific genes.

**Figure S7.**
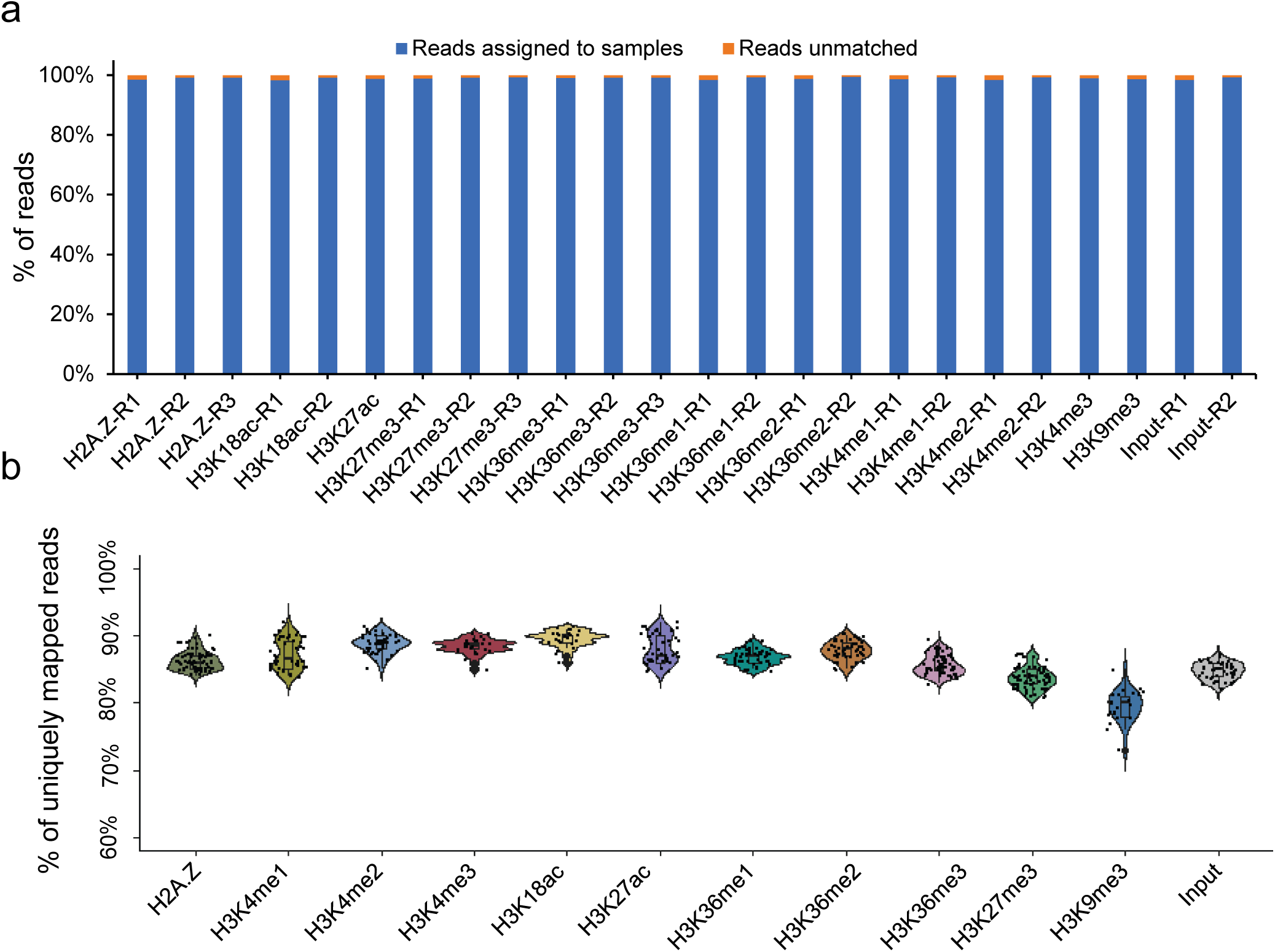
Pan-cancer multifactorial histone mark atlas profiling with mChIP-seq. a) Bar plot depicting de-multiplexing results of mChIP-seq data for 24 cancer cell lines. b) Violin plots depicting percentage of uniquely mapped reads of mChIP-seq data for 24 cancer cell lines.

**Figure S8.**
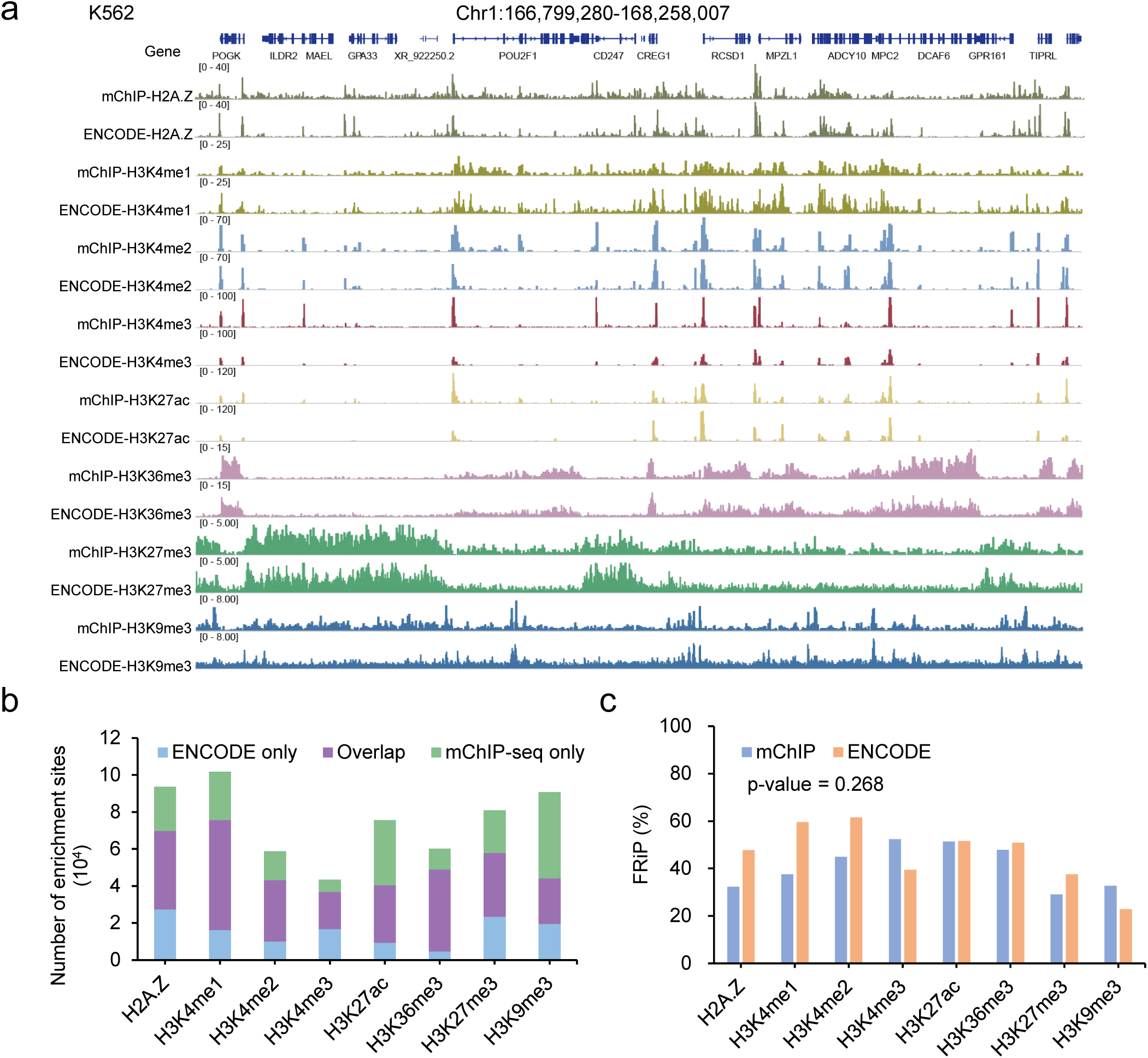
Performance of mChIP-seq for up to 24 samples in parallel. a) Genome browser tracks of K562 mChIP-seq and ENCODE data sets for comparison. b) Bar plot depicting enrichment sites shared between K562 mChIP-seq and ENCODE data sets. c) Bar plot comparing FRiP between mChIP-seq and ENCODE data sets from cell line K562. Merged peaks of each modification were used for calculating FRiPs. Paired t-test was used for p-value calculation.

**Figure S9.**
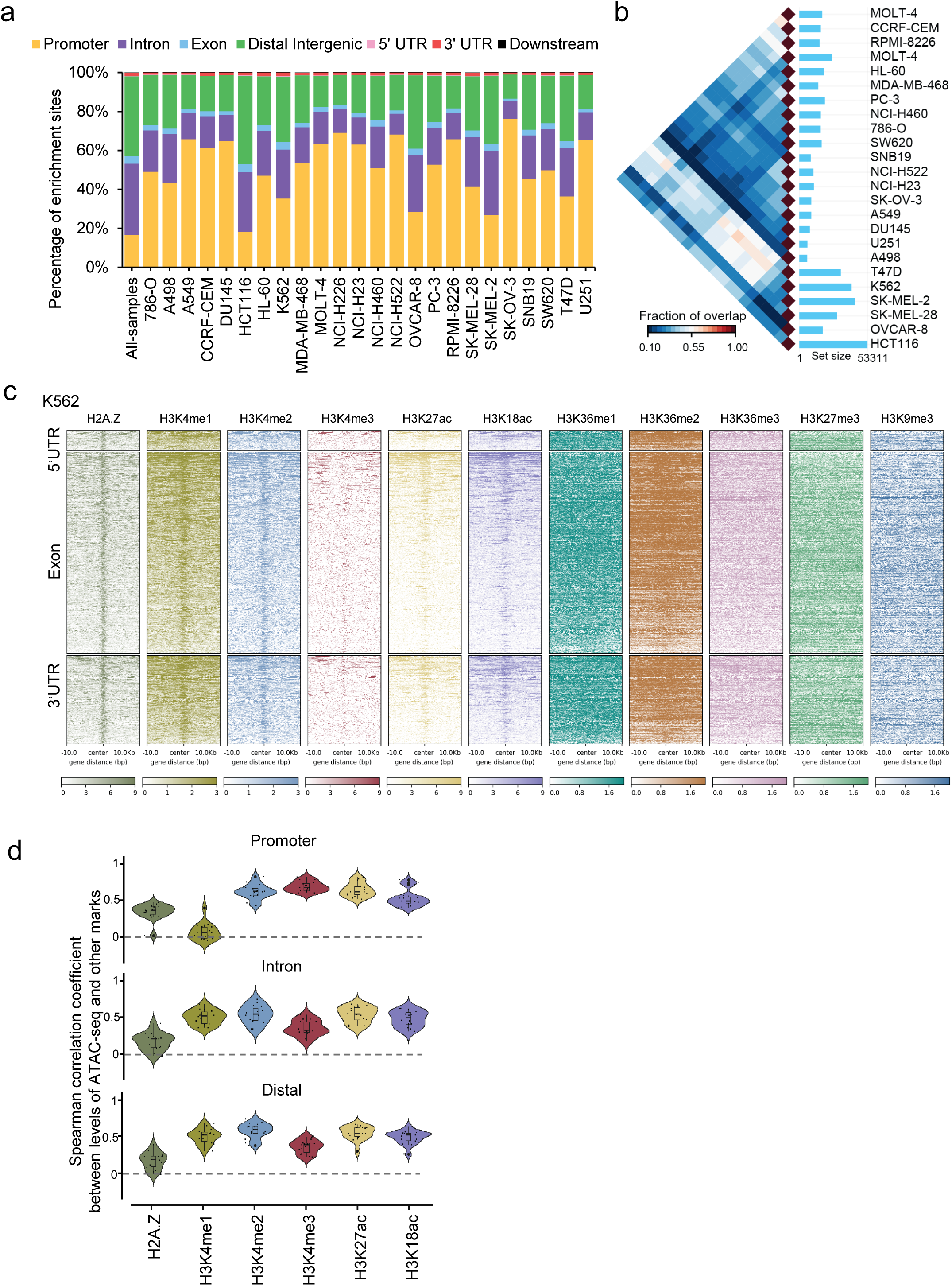
Epigenome characterization of H2A.Z in cancer cells. a) Percentage of H2A.Z enrichment sites overlapped to different types of the genomic regions. All-samples represents the merged peak set from all samples. Color indicates the type of genomic region overlapped by the peak. UTR: untranslated region. b) Pairwise intersection of enrichment sites for H2A.Z in the 24 cancer cell lines. c) Representative Heatmap showing signal intensities of 10 histone modifications around H2A.Z enrichment sites localized in 5’UTR, exon, and 3’UTR identified in K562. Signal intensities were sorted by H2A.Z data. d) Violin plots showing correlations of ATAC-seq signals to other histone marks for each cancer cell line in the H2A.Z enrichment sites localized in promoter, intron, and distal intergenic. 15 cancer cell lines with available ATAC-seq data were used for this analysis.

**Figure S10.**
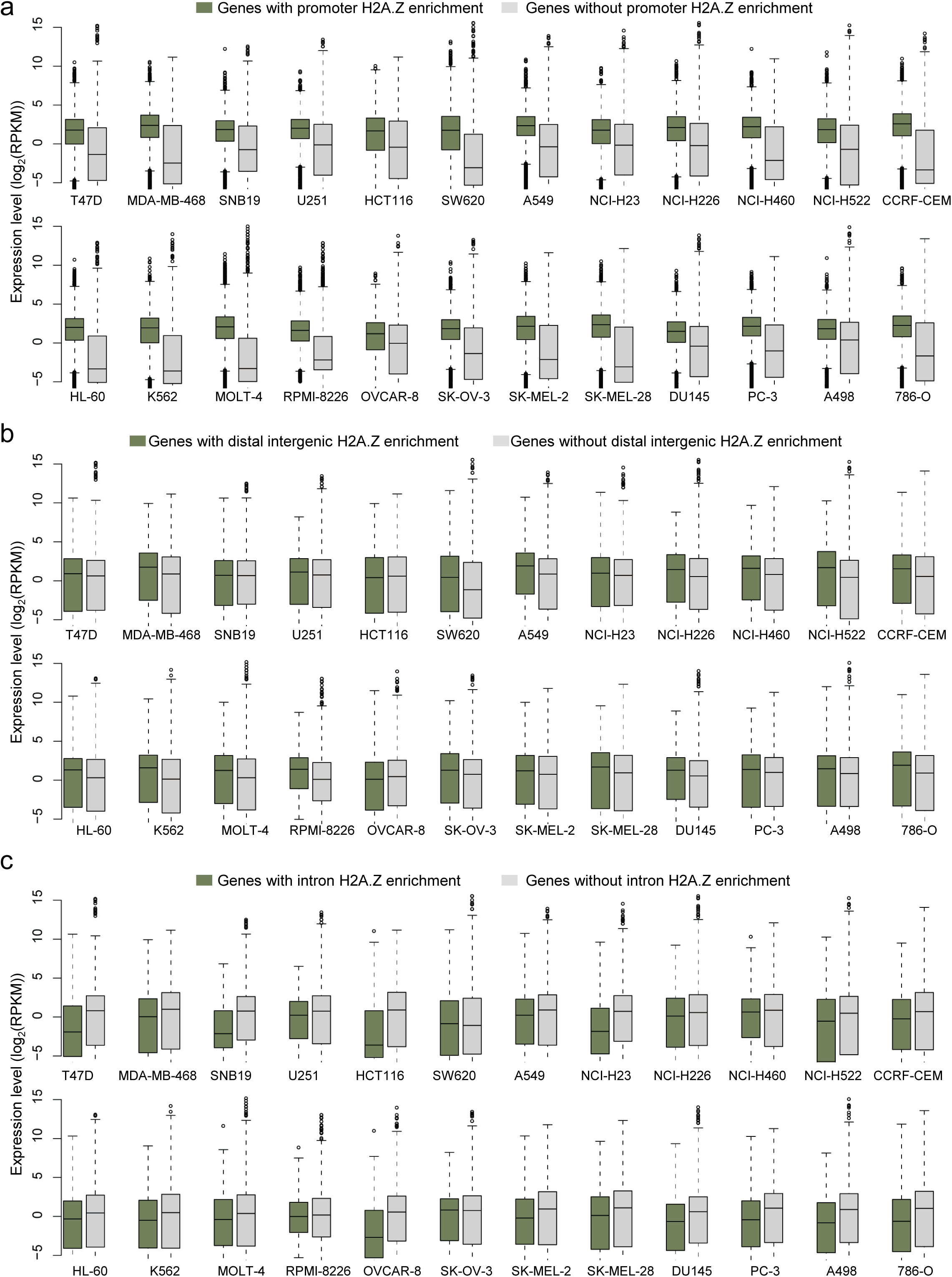
Associations of H2A.Z with gene expression in cancer cells. a) Boxplots comparing expression level of genes with promoter enrichment H2A.Z to genes without H2A.Z promoter enrichment. b) Boxplots comparing expression level of genes with intron H2A.Z enrichment to genes without H2A.Z intron enrichment. c) Boxplots comparing expression level of genes with distal intergenic H2A.Z enrichment to genes without H2A.Z distal intergenic enrichment.

**Figure S11.**
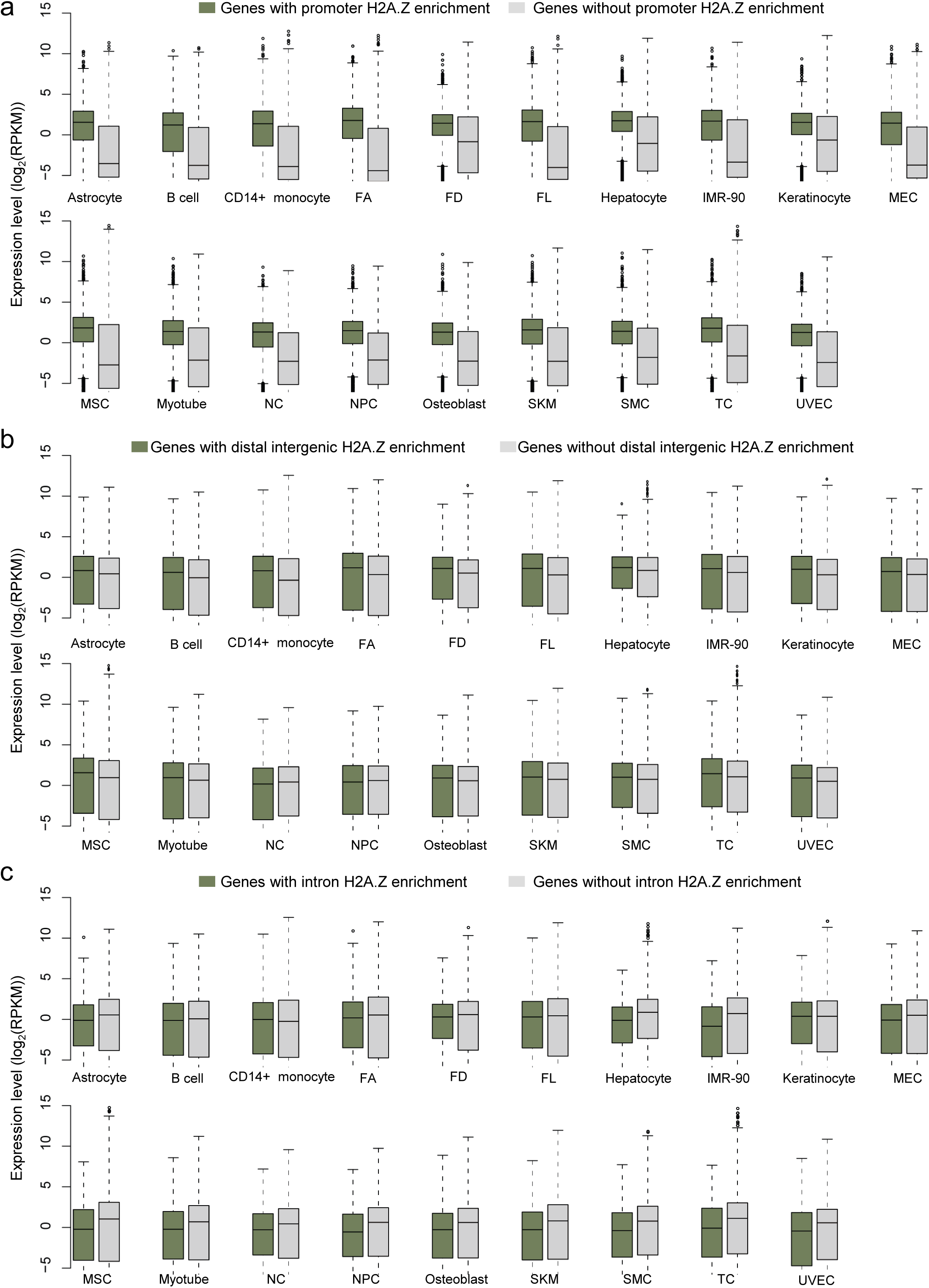
Associations of H2A.Z with gene expression in non-cancerous cells. a) Boxplots comparing expression level of genes with promoter enrichment H2A.Z to genes without H2A.Z promoter enrichment. b) Boxplots comparing expression level of genes with intron H2A.Z enrichment to genes without H2A.Z intron enrichment. c) Boxplots comparing expression level of genes with distal intergenic H2A.Z enrichment to genes without H2A.Z distal intergenic enrichment. FA: fibroblast of arm; FD: fibroblast of dermis; FL: fibroblast of lung; MEC: mammary epithelial cell; MSC: mesenchymal stem cell; NC: neural cell; NPC: neural progenitor cell; SKM: skeletal muscle myoblast; TC: trophoblast cell.

**Figure S12.**
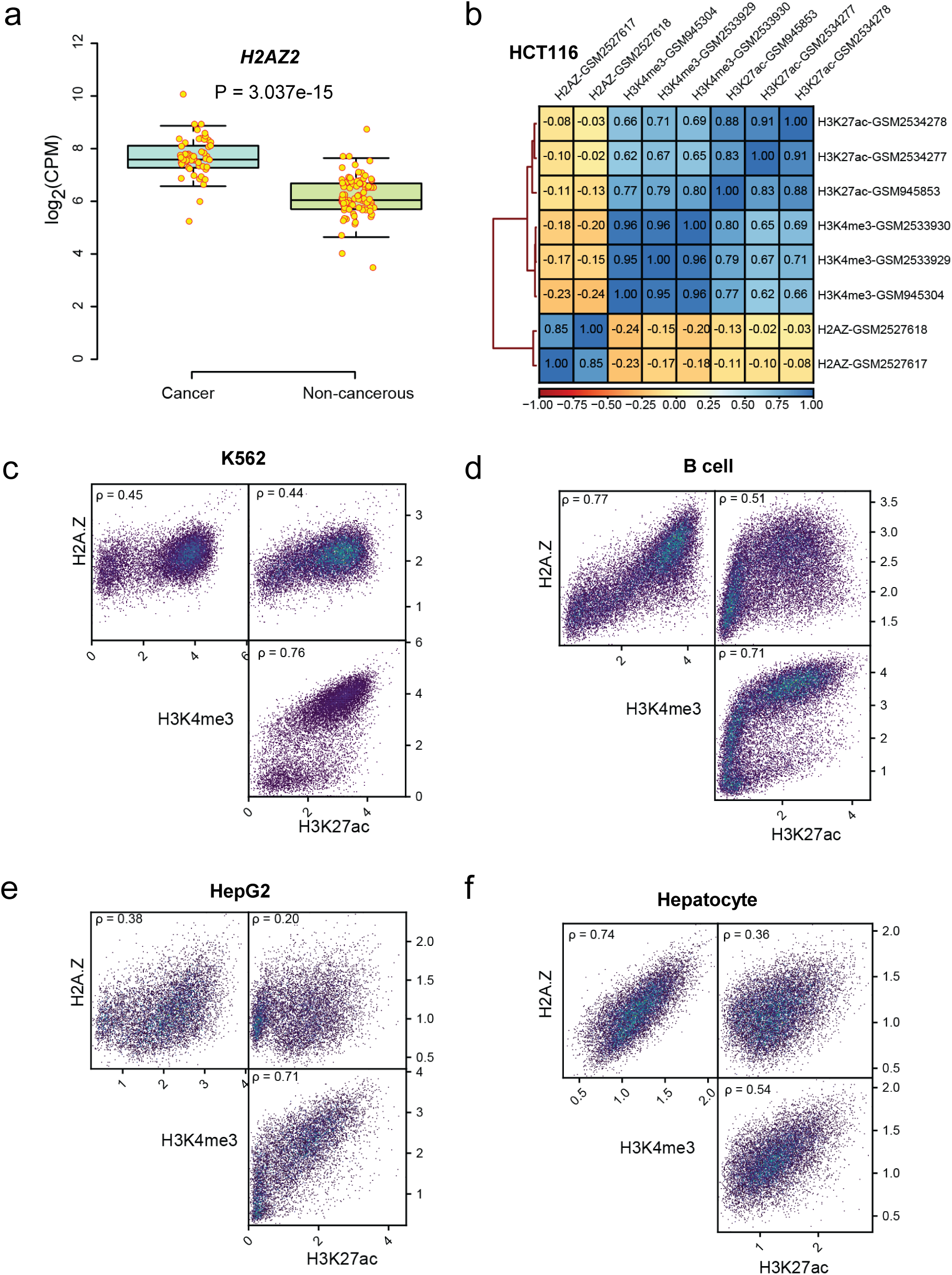
A decoupling of H2A.Z and H3K4me3 at promoter in cancer cells. a) Boxplot comparing expression of *H2AZ2* in cancer cells (n = 56) with non-cancerous cells (n = 65). P values were calculated using Mann-Whitney U tests. b) Heatmap showing signal correlations among H2A.Z, H3K4me3, and H3K27ac at promoter H2A.Z enrichment sites in HCT116 using data sets from ENCODE project. b) to f) Pairwise heatscatter plots showing the correlations of signal among H2A.Z, H3K4me3, and H3K27ac at promoter H2A.Z enrichment sites in K562 (c), B cell (d), HepG2 (e), and Hepatocyte (f). **ρ:** Spearman correlation coefficient.

**Figure S13.**
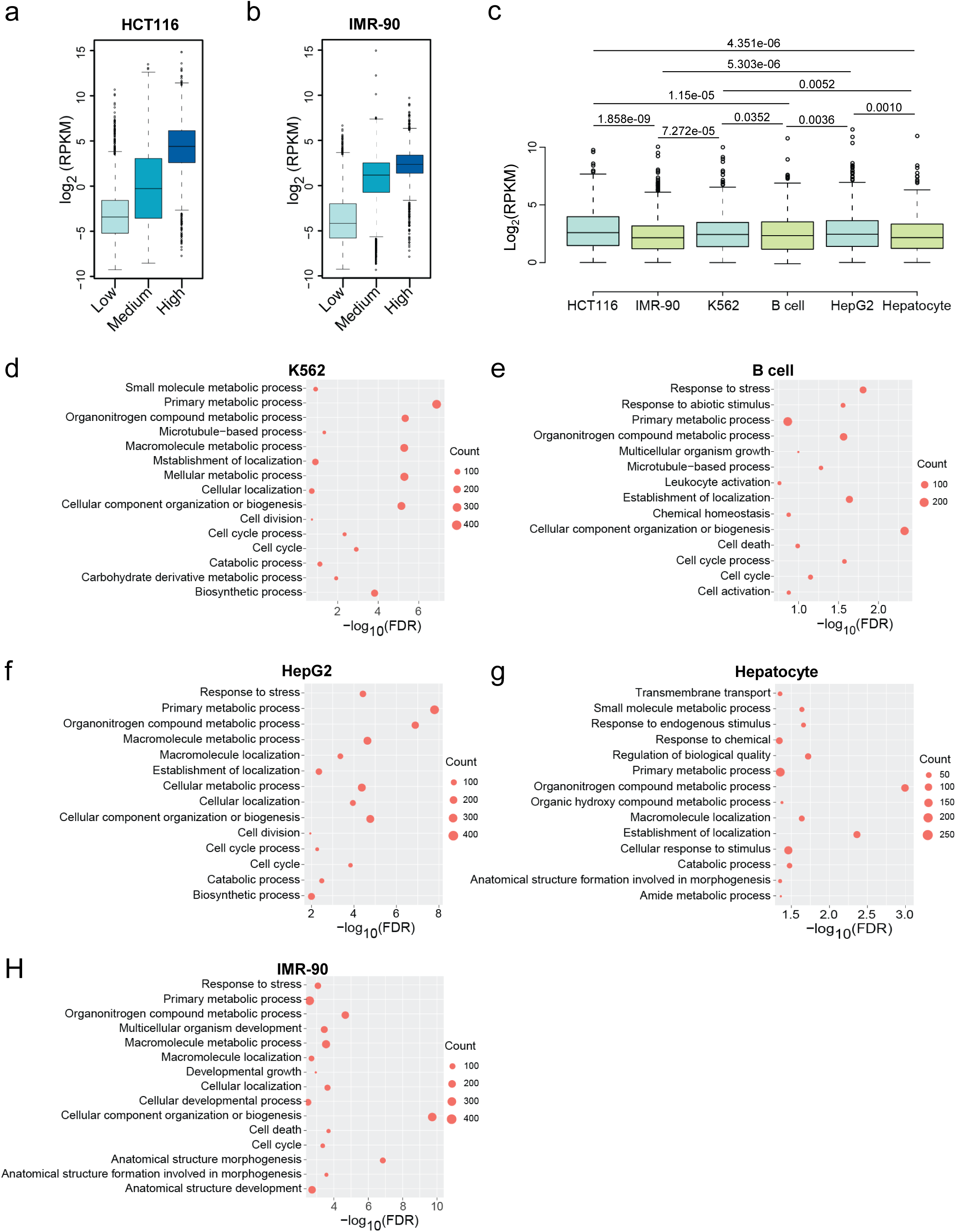
The impact of abnormal H2A.Z accumulation on gene expression at promoter regions in cancer cells. a) and b) Boxplots showing distribution of expression levels for genes associated with promoter H2A.Z enrichment sites proportionally divided into three groups by H3K4me3 intensities in HCT116 (a) and IMR-90 (b). P values calculated by Mann-Whitney U tests were < 2.2e-16 for all pairwise comparisons. c) Boxplot comparing levels of expressed genes (RPKM > 1) with H2A.Z (RPGC > 1) and without H3K4me3 (RPGC < 1) signals at promoter for cell lines HCT116, IMR-90, K562, B cell, HepG2, and Hepatocyte. P values were calculated using Mann-Whitney U tests. Only values of cancer cell lines compared to non-cancerous cell lines were presented. d) to h) Results of GO biological process enrichment analysis for expressed genes (RPKM > 1) with H2A.Z (RPGC > 1) and without H3K4me3 (RPGC < 1) signals at promoter in K562 (d), B cell (e), HepG2 (f), and Hepatocyte (g), and IMR-90 (h). Fifteen terms with the lowest FDR were presented.

**Figure S14.**
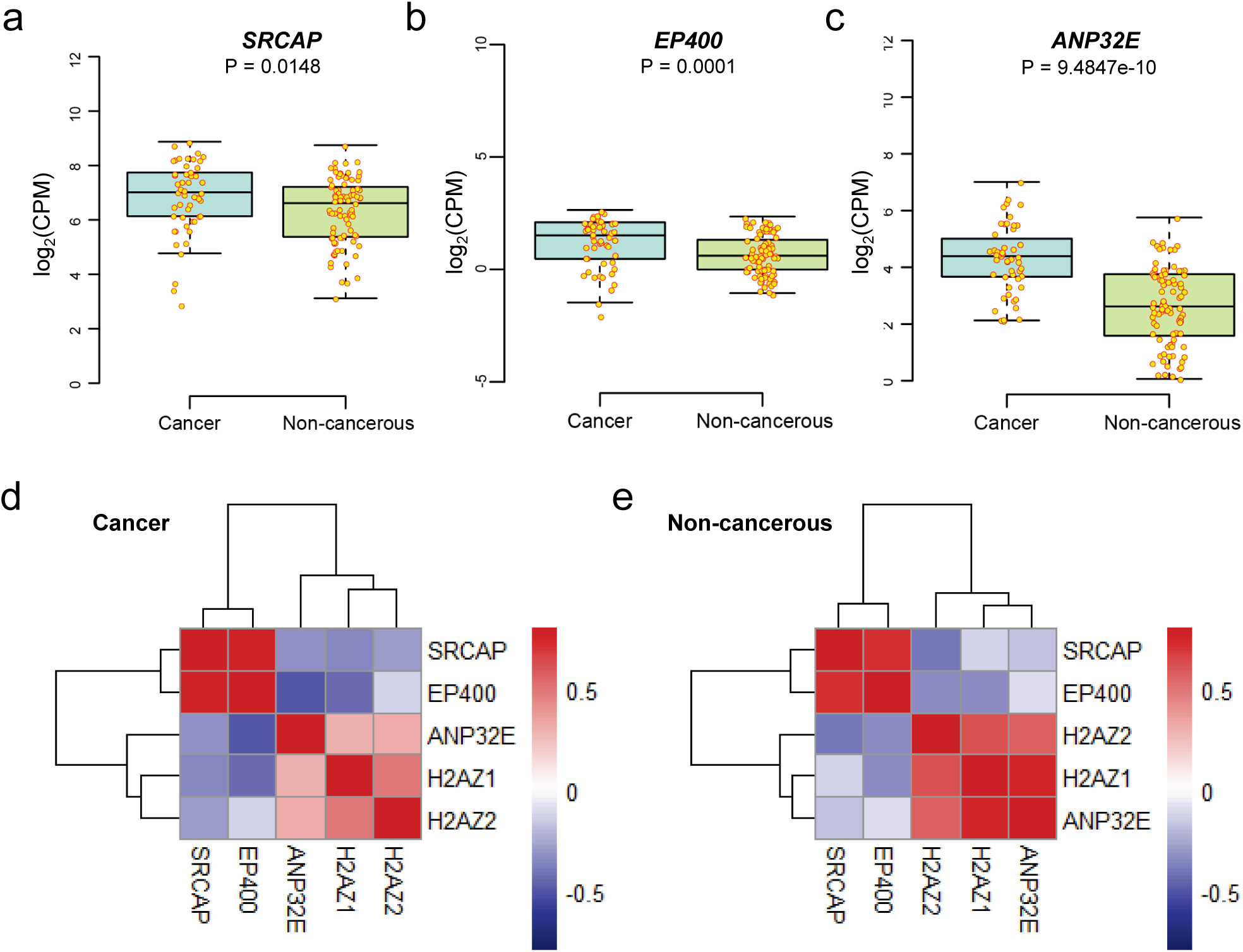
Expression analysis of chromatin remodelers involving in H2A.Z deposition. a) to c) Boxplot comparing expression of *SRCAP* (a)*, EP400* (b), and *ANP32E* (c) in cancer cells (n = 56) with non-cancerous cells (n = 65). P values were calculated using Mann-Whitney U tests. d) and e) Heatmap showing correlations of expression among *SRCAP, EP400, ANP32E, H2AZ1*, and *H2AZ2* in cancer cells (d) and non-cancerous cells (e).

**Figure S15.**
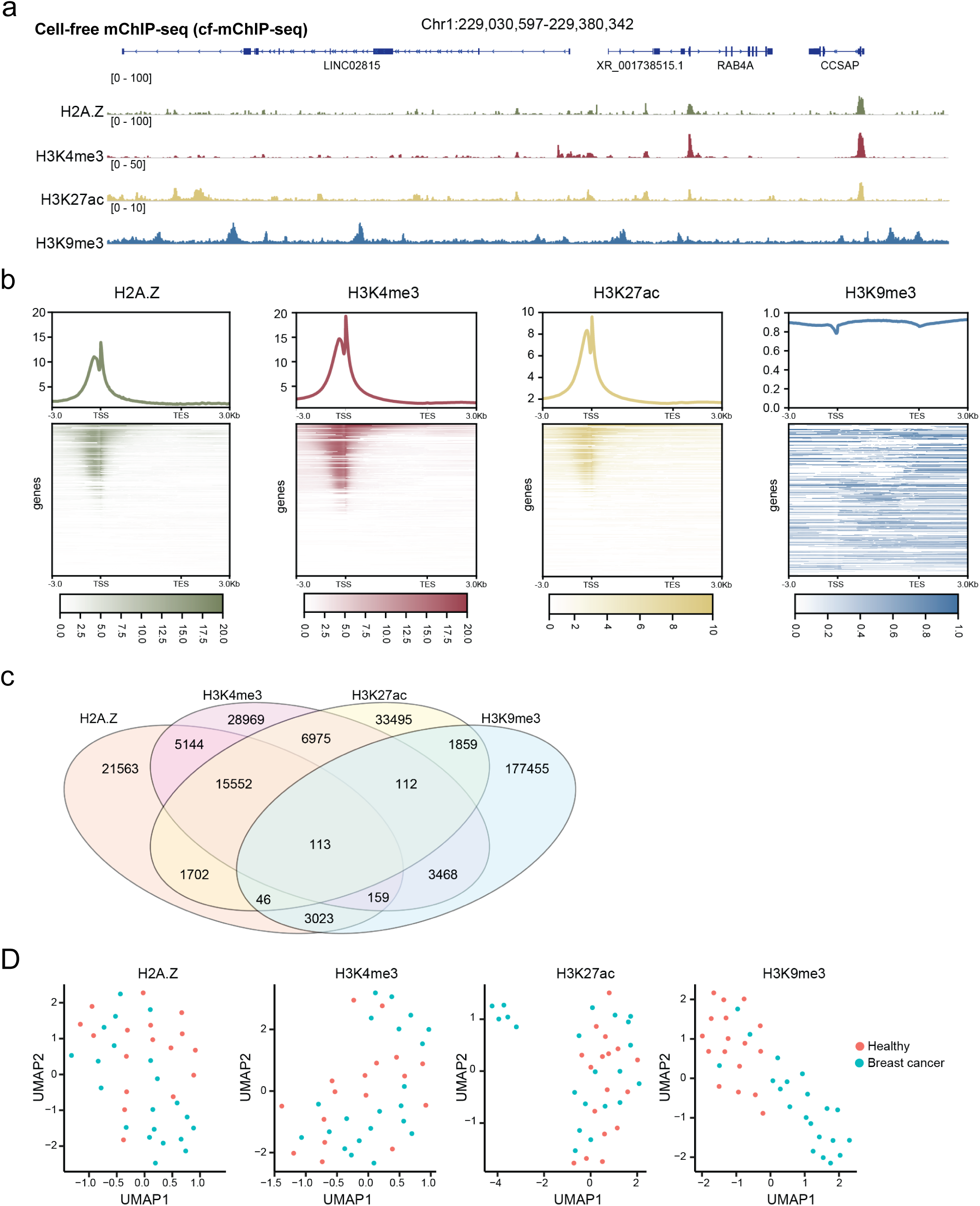
Epigenomic profiling of circulating nucleosomes in plasma from breast cancer patients using cf-mChIP-seq. a) Representative genome browser tracks of H2A.Z, H3K27ac, H3K4me3, and H3K9me3 signals in plasma profiled by cf-mChIP-seq. b) Plots (up) and heatmaps (down) showing normalized signal intensities of H2A.Z, H3K27ac, H3K4me3, and H3K9me3 in plasma over transcripts. Flanking regions are 3 kb upstream of TSSs and 3 kb downstream of the TESs. c) Venn diagram representing the number of peaks shared among H2A.Z, H3K27ac, H3K4me3, and H3K9me3 in plasma. d) Uniform manifold approximation projection (UMAP) plots showing plasma samples in two dimensions by using H2A.Z, H3K27ac, H3K4me3, H3K9me3 signals in peaks.

**Figure S16.**
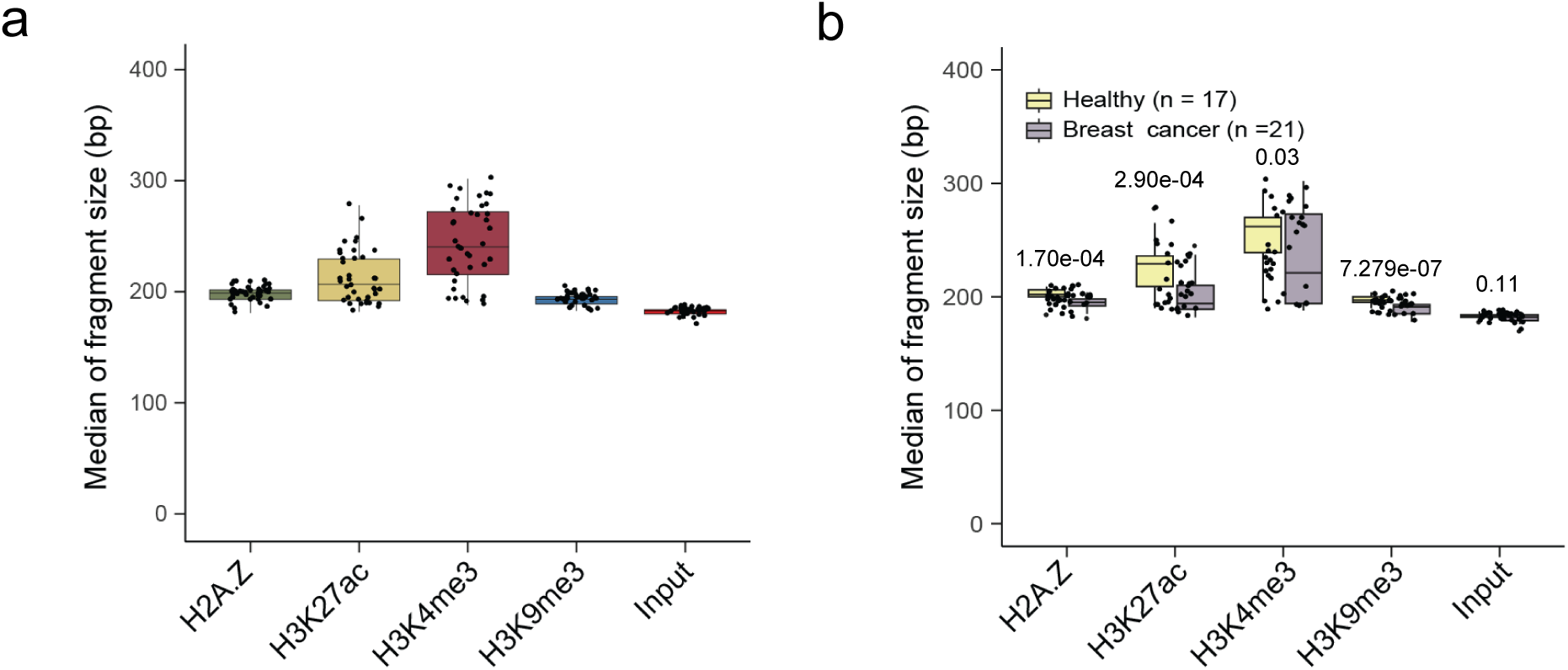
Fragment size of cfDNA associated with histone marks. a) Boxplot showing the median size of cfDNA fragments associated with histone marks H2A.Z, H3K27ac, H3K4me3, H3K9me3. Input (E) showed data without ChIP. Significant differences were observed when compared pairwise (P < 0.001, Paired Samples t-Test). b) Boxplot showing the difference of the median size of cfDNA fragments associated with histone marks H2A.Z, H3K27ac, H3K4me3, and H3K9me3 between patients with breast cancer (n = 21) and healthy controls. cancer status. P-values were calculated by using Welch’s t-test.

**Figure S17.**
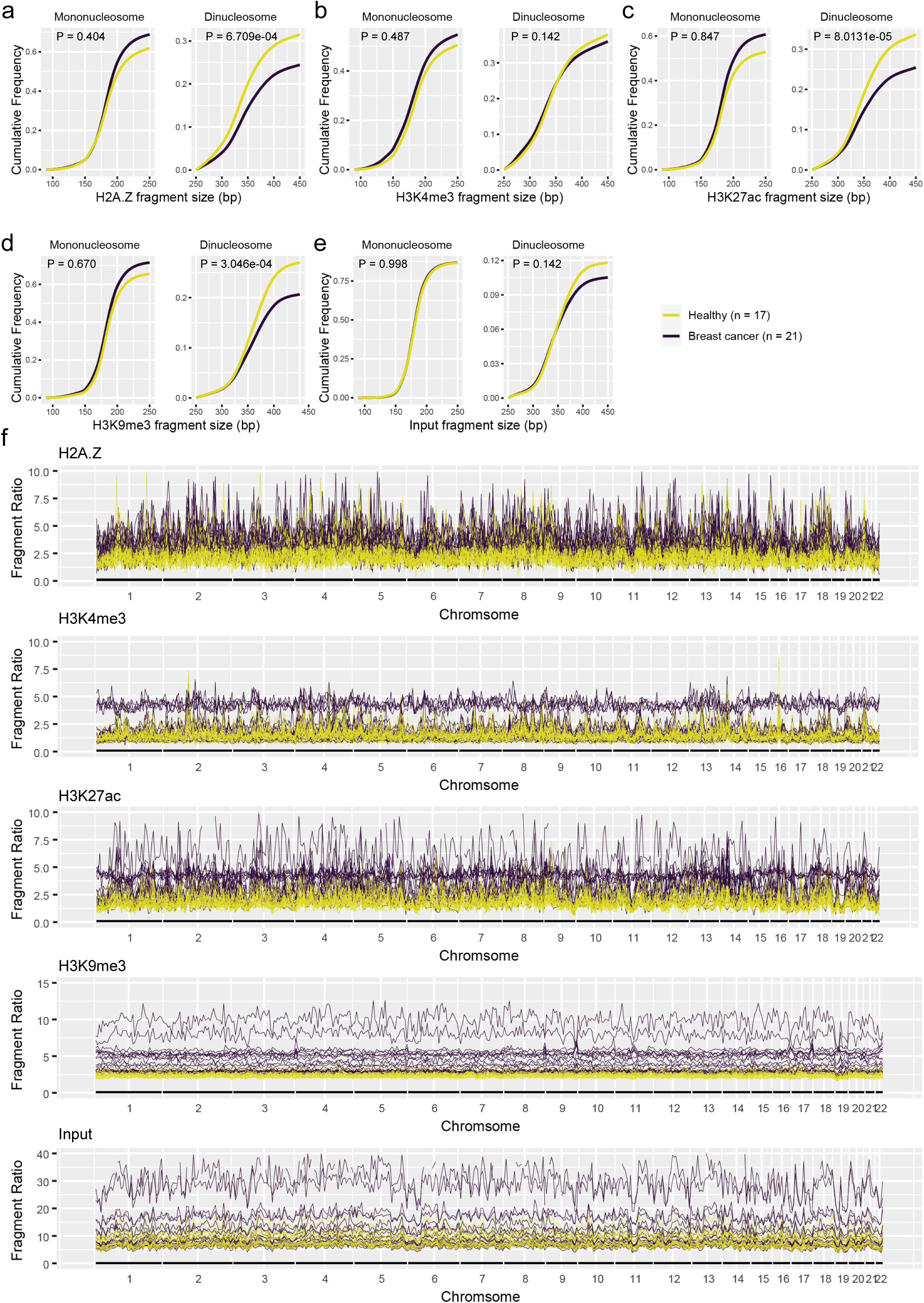
Fragmentation patterns of cfDNA associated with histone marks. a) to e) Line plots depicting the cumulative fragment size frequency normalized by total fragment counts across the mono- and di-nucleosome fractions of cfDNA associated with H2A.Z (a), H3K4me3 (b), H3K27ac (c), H3K9me3 (d) grouped by cancer status. Input (e) showed data without ChIP. f) The fragment ratio (the ratio of mono-nucleosome counts to di-nucleosome counts) showing genome-wide fragmentation patterns of histone-mark-associated cfDNA profiles in 5-Mb bins for each sample.

### Supplementary Tables

**Table S1. mChIP-seq compared to existing high-throughput epigenomic profiling methods.**

**Table S2. Information of libraries generated by mChIP-seq for 24 cancer cell lines.**

**Table S3. Information of data from public database used in this study.**

**Table S4. Information of libraries generated by cf-mChIP-seq. Table S5. List of key resources used in this study.**

## Notes

### Competing Interest Statement

The authors have declared no competing interest.

